# Claspin is required for growth recovery from serum starvation through regulating the PI3K-PDK1-mTOR pathway

**DOI:** 10.1101/2022.01.21.475743

**Authors:** Chi-Chun Yang, Hisao Masai

**Affiliations:** Department of Basic Medical Sciences, Tokyo Metropolitan Institute of Medical Science, 4-6-1 Kamikitazawa, Setagaya-ku, Tokyo 156-8506, Japan

**Author notes:** Correspondence should be addressed to Hisao Masai.

**Keywords:** Claspin, serum stimulation, PI3 kinase, PDK1, mTOR

## Abstract

Growth recovery from serum starvation requires the activation of PI3 kinase (PI3K)-PDK1-Akt-mTOR pathways. Claspin plays multiple important roles in regulation of DNA replication as a mediator for the cellular response to replication stress, an integral replication fork factor that facilitates replication fork progression and a factor that promotes initiation by recruiting Cdc7 kinase. Here, we report a novel role of Claspin in growth recovery from serum starvation. In the absence of Claspin, cells do not proceed into S phase and eventually die. Claspin interacts with PI3K and mTOR, and is required for activation of PI3K-PDK1-mTOR and for that of mTOR downstream factors, p70S6K and 4E-BP1, but not for p38 MAPK cascade during the recovery from serum starvation. PDK1 interacts with Claspin, notably with CKBD, in a manner dependent on phosphorylation of the latter protein, and is required for interaction of mTOR with Claspin. p53 and ROS (Reactive Oxygen Species) inhibitors increased survival of Claspin-deficient cells released from serum starvation. Thus, Claspin plays a novel role as a mediator/protein platform for nutrition-induced proliferation/survival signaling by activating the mTOR pathway.

## Introduction

Living organisms are exposed to various types of stresses, but they have so called “stress response systems” to cope with these situations. For example, in the cellular response pathway to replication stress, a signal is transmitted from sensor kinase (ATR) to effector kinase (Chk1) to temporarily arrest progression of replication and cell division. Claspin functions as a mediator of ATR to Chk1 signaling in the replication stress response (Branzei and Foiani, 2009). The stalled replication fork caused by replication stress results in activation of the ATR-ATRIP complex and activated ATR phosphorylates Chk1 at Ser317 and Ser345 to block DNA replication (Zhao, H. and Piwnica-Worms, H., 2001). Chk1 phosphorylation by ATR requires Claspin as a mediator for the checkpoint signal transduction. ATR phosphorylates Chk1 in the Claspin-Chk1 complex; this phosphorylation occurs more efficiently *in vitro* on the Claspin-Chk1 complex than on the Chk1 alone (Lindsey-Boltz et al., 2009).

Claspin was originally discovered as a factor that binds to Chk1 and promotes replication checkpoint in *Xenopus* egg extract (Kumagai and Dunphy, 2000). Mrc1, its yeast homolog, is also essential for activation of downstream effector kinases (Rad53 and Cds1/Rad53, respectively, in budding and fission yeasts). Thus, the role of Claspin/Mrc1 as a replication checkpoint mediator is well conserved from yeasts to human (Tanaka and Russell, 2001; Alcasabas et al., 2001; Chini and Chen, 2003; Osborn and Elledge, 2003; Yoo et al., 2006; Lindsey-Boltz et al., 2009). The Chk1 binding domain of Claspin (CKBD) binds Chk1 in a manner strictly dependent on its phosphorylation (Kumagai and Dunphy, 2003). Thr-916 residue in CKBD is phosphorylated after HU treatment and Chk1 cannot be activated in human cells carrying the T916A/S945A mutation in Claspin (Chini and Chen, 2006). It was reported that Cdc7 kinase and casein kinase1 γ1 are potential kinases for phosphorylating CKBD under DNA replication stress condition (Meng et al., 2011; Yang et al., 2019).

In addition to the Claspin’s role as a checkpoint adaptor, Claspin may also play important roles in regulation of normal course of DNA replication, similar to Mrc1 (Osborn AJ & Elledge SJ, 2003; Szyjka SJ, et al., 2005; Tourriere H, et al., 2005; Yeeles, et al., 2017; Petermann, et al.,2008; Yang, et al., 2016). Claspin has been shown to be required for efficient DNA replication through its ability to facilitate the replication fork movement (Petermann, et al., 2008). We have recently reported a novel function of Claspin in replication initiation control; Claspin promotes phosphorylation of Mcm by recruiting Cdc7, the kinase essential for initiation of DNA replication.

Recent reports show that Mrc1 in budding yeast, the ortholog of Claspin, plays a role in slowing down DNA replication in response to osmotic, oxidative, heat and nutrient deficiency stresses (Duch A, et al., 2013; Duch A, et al., 2018). In mammalian cells, Claspin is required for suppression of replication origin firing and for slowing DNA replication fork progression in response to endoplasmic reticulum stress (Nadal et al., 2015, Cabrera et al., 2017). These cellular pathways are independent of the replication checkpoint pathways, indicating roles of Mrc1/Claspin in stress response pathways other than replication checkpoint.

Growth factors, like PDGF, FGF and IL3, have been reported to promote cell survival and cell cycle progression (McDonald and Hendrickson, 1993; (Plotnikov, et al., 2000; Stauber, et al., 2000; Chang, F., et al., 2003). Interactions of these factors with their receptors trigger the activation of phosphoinositide 3-kinase (PI3K) (Toker, A., and L. C. Cantley. 1997; Vanhaesebroeck, et al., 1997). PI3K recruits Phosphoinositide-dependent protein kinase 1 (PDK1), leading to activation of Akt and mTOR (Williams et al., 2000; Toker and Newton, 2000; Belham C et al. 1999). Mechanistic target of rapamycin (mTOR), an atypical serine/threonine kinase, belongs to the phosphatidylinositol 3-kinase-related family (PIKK). mTORC could accept many intracellular and extracellular signals, such as amino acids, growth factors, energy, oxygen, and DNA damage. Nutrient sensing or responses to nutrient utilization are among of the main functions of the mTORC1, and the signal transmission originating from the levels and quality of nutrients requires the coordinated interaction of sensors and regulators upstream of mTORC1 (Ma et al., 2018). When sufficient nutrients are available, mTOR is phosphorylated at Ser2448 via the PI3K/Akt signaling pathway and transmits signals to downstream factors p70 S6 kinase and eIF4E inhibitor 4E-BP1 to promote cell growth and cell cycle progression (Navé et al., 1999; Fang Y, et al., 2001; Pullen et al., 1997; Dufner et al., 1999). However, potential roles of Claspin in the nutrient-induced PI3K-PDK1-mTOR pathway have not been explored.

In this study, we show that Claspin is required for growth restart from serum starvation-induced growth arrest, and interacts with the ATR-like PIKK, PI3K, and factors in the mTOR pathway in a manner independent of the ATR-Chk1 pathway. Claspin interacts with PI3K, PDK1 and mTOR and promotes phosphorylation of factors involved in the PI3K-PDK1-mTOR pathway, when cells are released from serum starvation. The findings in this study show a potential role of Claspin as a protein hub for growth-dependent activation of the PI3K-PDK1-mTOR pathway.

## Results

### Growth recovery from serum starvation requires Claspin

We previously established mouse embryonic fibroblast cells in which Claspin can be inducibly knocked out. In this cell line, the exon 2 are flanked by two loxP sites, and can be excised out by expressing Cre recombinase, leading to knockout of the Claspin gene. We noted that during serum starvation, parts of Claspin knockout MEF cells appeared apoptotic. Upon Claspin knockout, Annexin V-positive cells in an early apoptotic stage increased from 5 % to 42.9 % at 48 hr after serum starvation. In TUNEL assays, cells that contain fragmented DNA increased from 10.8 % to 64.5 % at 48 hr after serum starvation. However, analyses of DNA content indicated the sub-G1 population increased only mildly (from 5.3% to 8.1%), indicating that they did not progress into an incorrigible stage (**Supplementary Fig. S1A and B**). However, during the growth recovery from the serum starvation, MEF cells lacking the Claspin gene underwent massive cell death (**Fig. 1A and B**). Furthermore, 293T cells depleted of Claspin by siRNA also similarly died after release from serum starvation (**Supplementary Fig. S2A and B**). We examined other replication and checkpoint-related factors for their roles in growth restart after serum starvation. Knockdown of Cdc7 and CKγ1, but not that of Tim, AND1, TopBP1 and Cdc45, caused cell death during serum-induced growth in 293T cells (**Supplementary Fig. S2C**). This indicates that Claspin’s roles in growth from serum starvation is not simply due to its role in replication checkpoint or to that in replication initiation. We also noted that some population of the surviving Claspin knockout cells became senescent (**Supplementary Fig. S3A and B**).

**Fig. 1.**
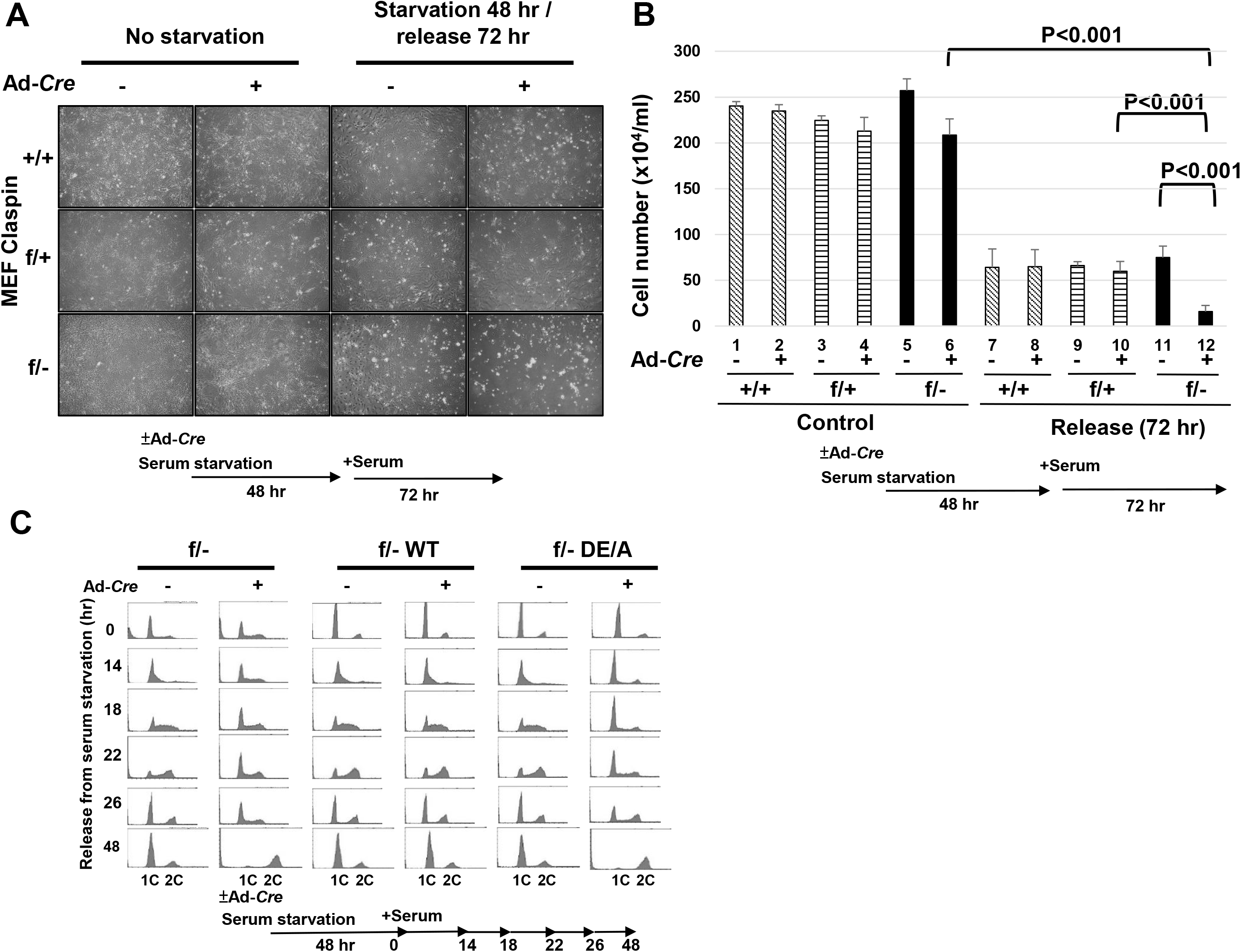
Claspin is required for growth recovery from serum starvation. (**A**) *claspin(+/+)*, *(f/+)* and *(f/-)* MEF cells, treated with Ad-*Cre* or untreated, were cultured in serum-free medium for 48 hr. After serum was added to the medium, the cells were grown for 72 hr and observed under microscopy. (**B**) Quantification of the cell numbers of the cultures in (**A**). Experiments were repeated 3 times and error bars are indicated. (**C**) *claspin (f/-)* MEF cells and the same cells carrying claspin transgene (WT, DE/A), treated with Ad-*Cre* or untreated, were cultured in serum-free medium for 48 hr. After addition of serum at time=0, cells, harvested at indicated times, were analyzed by FACS.

We previously identified an acidic patch segment (AP) on Claspin which is required for replication initiation and replication stress functions through its ability to recruit Cdc7 kinase. We examined whether AP is required for growth recovery from serum. We ectopically expressed the wild-type and DE/A mutant of Claspin in Claspin (f/-) MEF cells. The DE/A mutant, in which all the acidic residues in AP were replaced with alanine, cannot interact with Cdc7. We analyzed the DNA contents of the cells after release from serum starvation by flow cytometry (**Fig. 1C**). The control cells entered S phase by 14 hr after release, and nearly completed it by 22 hr, and returned to G1 by 26 hr. On the other hand, the Claspin knockout cells did not enter S phase or only very slowly replicated its genome by 26 hr, and at 48 hr, most surviving cells were found to have G2 DNA content. Exogenous expression of the wild-type Claspin restored the timely S phase entry and reversed the cell death effect in the Claspin knockout cells. In contrast, the DE/A mutant could not rescue the defects, indicating that AP is essential for growth restart from serum starvation. Thus, AP is specifically required for entry into S phase during regrowth from serum starvation-induced G0 state.

### Effect of Claspin knockout on replication and cell cycle-related factors

Delayed S phase entry in Claspin knockout may be related to its role in initiation and elongation of DNA synthesis. Therefore, we have examined the factors involved in the process of DNA replication and cell cycle progression during the release from serum starvation. Phosphorylation of MCM (MCM2 (S53) and MCM4 (S6T7)) was induced by 14 hr in the control, but was not induced in Claspin knockout, consistent with the fact that cells do not enter S phase in the absence of Claspin (**Fig. 2A and C**). Histone H3 S10 phosphorylation increased from 18 to 48 hr in control, consistent with active cell division, but it did not increase after 22 hr in Claspin knockout, consistent with failure of cell division (**Fig. 2B**). Activation of MAP kinases, p38 and ERK1/2, observed during growth stimulation, is not significantly affected by loss of Claspin (**Fig. 2A**). Rb protein increased after growth stimulation in control, but did not increase in Claspin knockout, whereas the total p53 protein as well as the activated form of p53 increased in Claspin knockout, and stayed at the levels higher than that found in the control (**Fig. 2D and 2E**). The γH2AX signal moderately increased after release from serum starvation in Claspin knockout cell, indicating the presence of DSB (**Fig 2B**). However, Chk1 was not activated during serum starvation as well as during release from serum starvation in MEF cells (**Fig. 2A** **and Supplementary Fig. S4**), indicating that replication checkpoint is not activated.

**Fig. 2.**
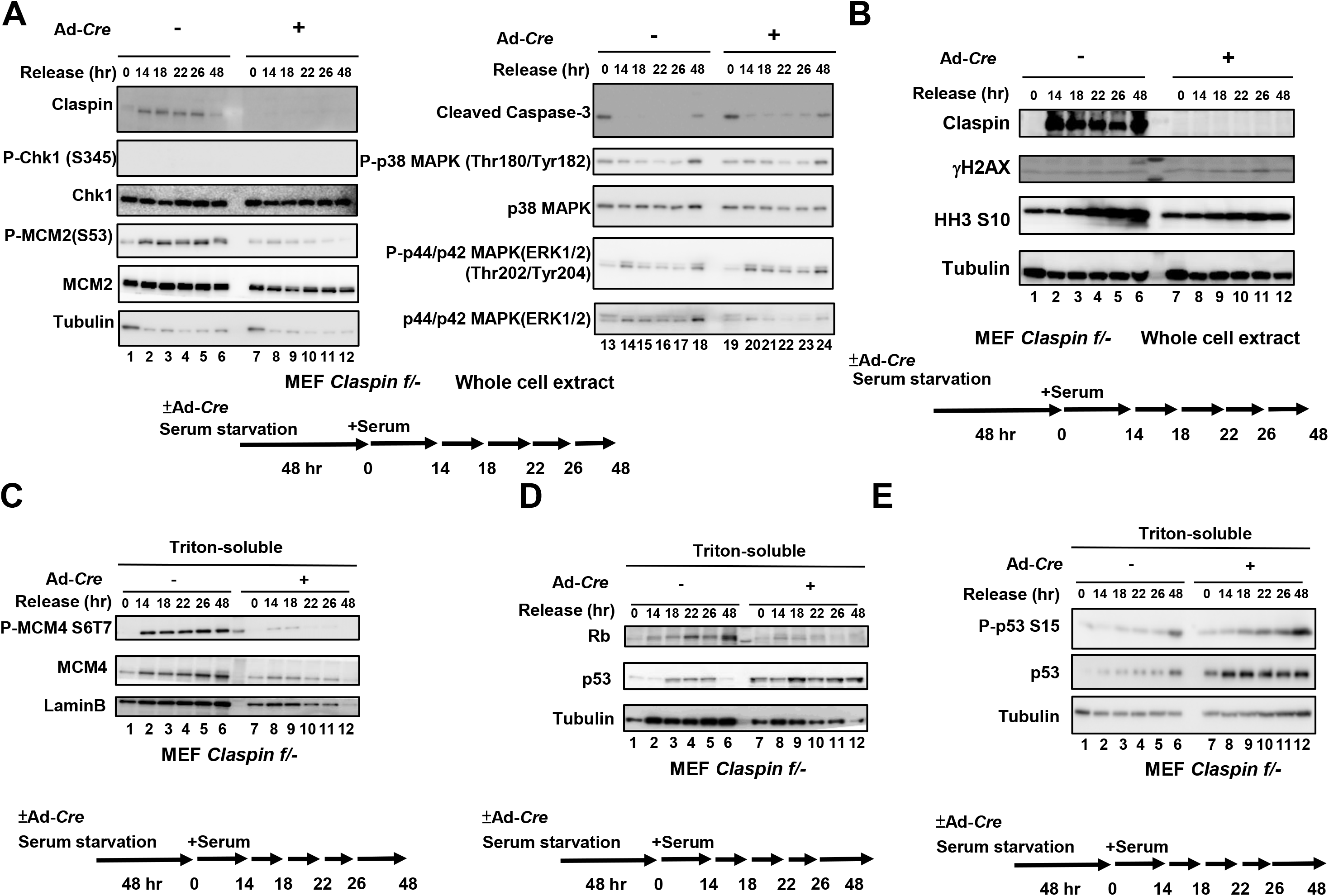
Effects of Claspin knockout on factors involved in DNA replication, cell cycle progression and MAP kinase pathways during the growth recovery from serum starvation. (**A**, **B**, **C**, **D** and **E**) *claspin (f/-)* MEF cells, treated with Ad-*Cre* or untreated, were cultured in serum-free medium for 48 hr. After addition of serum, cells were harvested at the indicated times and proteins were analyzed by western blotting with indicated antibodies. (**A** and **B**) Whole cell extracts were analyzed. (**C**, **D** and **E**) Cells were fractionated into Triton-soluble and -insoluble fractions, and Triton-soluble fractions were analyzed by western blotting with indicated antibodies.

### Claspin is required for activation of the PI3K/PDK1/mTOR pathway

The PI3K/PDK1/mTOR pathway is intimately related to the nutrient-mediated regulation of cell growth (Jewell and Guan, 2013). Therefore, we examined the effect of Claspin knockout on this pathway. Expression of PI3 kinase ClassIII is maintained after release for 26 hr and increased during the next 22 hr. On the other hand, it increased during the first 14 hr but decreased after that in Claspin knockout cells. Tyr 458 phosphorylation of PI3 kinase p85 subunit was observed at 48 hr in control, but was not detected in the Claspin knockout (**Fig. 3**). S473 phosphorylation of Akt (phosphorylated by mTORC2 and required for its activation) continues to increase after release until 48 hr, but that in Claspin knockout slightly increased at 14-18 hr and then decreased (**Fig. 3**). S2448 phosphorylation in mTOR, an indicator of mTOR activation, increased after release and maintained its expression until 48 hr after release, but that in Claspin knockout stayed low after release and started to decrease at 18 hr (**Fig. 3**). Phosphorylation of T37/46 of 4EBP1 (a target of mTORC1), which continued to increase in the control, also behaved in a similar manner in Claspin knockout, decreasing at 18 hr and thereafter (**Fig. 3**). On the other hand, phosphorylation of S241 of PDK1 (an indicator of active PDK1) increased until 48 hr in the wild-type, while it stayed high in the first 22 hr and then gradually decreased thereafter in Claspin knockout. In contrast, phosphorylation of S6K T389 was not activated at all in Claspin knockout, whereas it continued to increase until 48 hr in control (**Fig. 3**). These results show that the PI3K/PDK1/mTOR pathway is compromised in Claspin knockout cells during the growth recovery from serum starvation. Thus, Claspin is required for sustained activation of the PI3K/PDK1/mTOR pathway but not for growth-induced MAP kinase pathway during growth stimulation from the quiescent state.

**Fig. 3.**
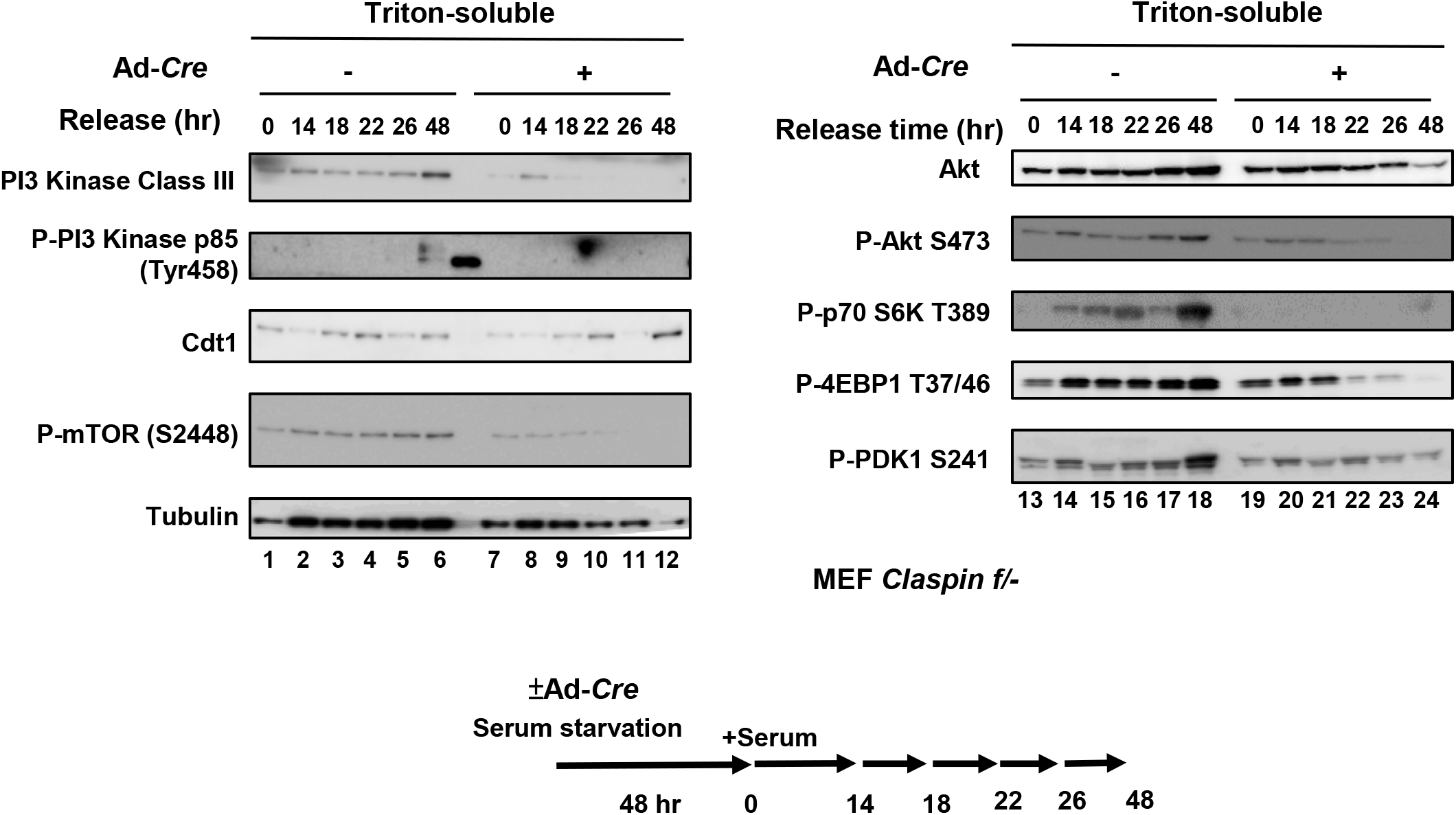
Effects of Claspin knockout on factors involved in the PI3K-Akt-mTOR pathway during the growth recovery from serum starvation. *claspin (f/-)* MEF cells, treated with Ad-*Cre* or untreated, were cultured in serum-free medium for 48 hr. Cells, harvested at the indicated times after addition of serum, were fractionated into Triton-soluble and -insoluble fractions (using 0.3% Triton X-100) and Triton-soluble fractions were analyzed by western blotting with indicated antibodies.

During serum starvation in 293T cells, the levels of phosphorylated PDK1 and mTOR were not affected by Claspin depletion, whereas they decreased in Claspin-depleted cells during the release from serum starvation, as was observed in Claspin knockout MEF cells (**Supplementary Fig. S5**). Chk1 was slightly activated during serum starvation, but not activated during the release. Exogenous expression of the DE/A mutant in Claspin knockout MEF cells did not rescue the phosphorylation of PDK1, mTOR, S6K or 4EBP1, indicating that the acidic patch is essential for activation of PI3K/PDK1/mTOR (**Supplementary Fig. S6 and** **Fig. 1C**).

### Cell death is induced by loss of Claspin during growth restart from serum starvation and is partially rescued by inhibition of p53 or ROS

Most cells died by 48 hr after growth stimulation in the absence of Claspin, as indicated by the FACS analyses (**Fig. 1A** and **C**). The p53 level increases and activated during this process (**Fig. 2E**), and indeed, addition of Pifithrin-α, a p53-specific inhibitor, at 50 µM partially rescued the cell death (**Fig. 4A**), indicating that the cell death at least partly depends on p53.

**Fig. 4.**
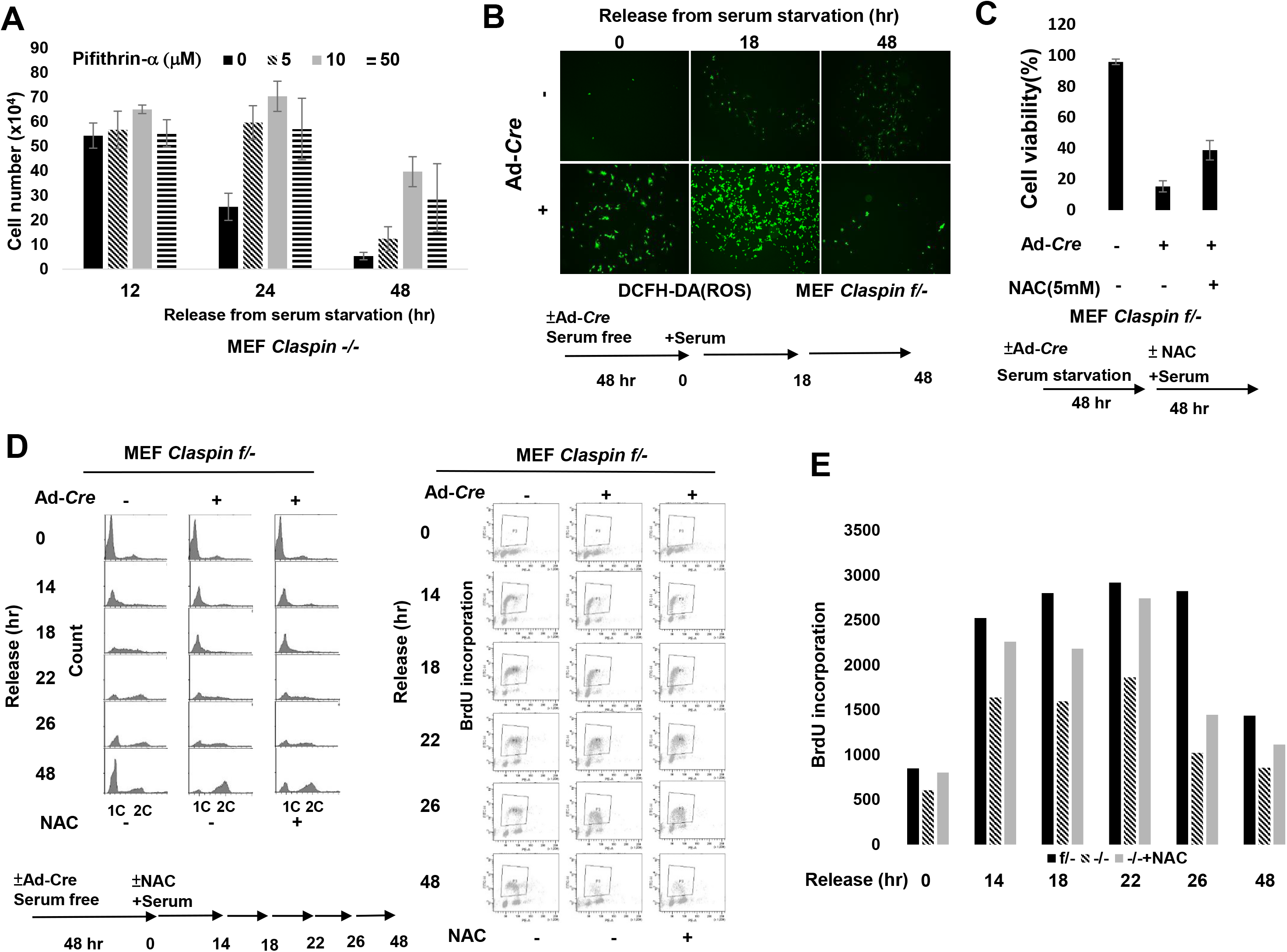
Roles of p53 and ROS in cell death and DNA synthesis during the growth recovery from serum starvation. (**A**) *claspin (f/-)* MEF cells, treated with Ad-*Cre*, were cultured in serum-free medium for 48 hr. After addition of serum and Pifithrin-α, a p53-inhibitor, at various concentrations, cells were harvested at the indicated times and observed under microscope. Viable cell numbers were counted and quantified. Experiments were repeated 3 times and error bars are indicated. (**B**) *claspin (f/-)* MEF cells, treated with Ad-*Cre* or untreated, were cultured in serum-free medium for 48 hr. After addition of serum, cells were treated with DCFH-DA (2’, 7’-Dichlorodihydrofluorescin diacetate; ROS indicator) at indicated times and then observed under florescence microscopy. (**C**) *claspin (f/-)* MEF cells, treated with Ad-*Cre* or untreated, were cultured in serum-free medium for 48 hr. Serum was added in the presence or absence of 5 mM NAC, and observed under microscope at 48 hr after release. Fractions of live cells were calculated and presented. (**D**) *claspin (f/-)* MEF cells, treated with Ad-*Cre* or untreated, were cultured in serum-free medium for 48 hr. At t=0, serum was added with or without NAC, and BrdU was added to the cells for 30 min at the indicated times. DNA contents were analyzed by FACS. (**E**) Quantification of BrdU incorporation from the data in (**D**).

ROS (Reactive Oxygen Species) often becomes a primary cause for genomic instability. The mTORC1 repressors such as Sesn1, TSC2, AMPKβ1, and IGF-BP3 also play roles in the regulation of ROS (Budanov AV & Karin M, 2008; Feng Z, et al,. 2007). p53 activates these mTORC1 repressors, resulting in lower metabolic activity and increased oxidative stress (A.V. Budanov et al,. 2004). Therefore, we examined production of ROS after release from serum starvation. In Claspin knockout MEF cells, more ROS were produced after serum starvation, and the level of ROS further increased at 18 hr after release (**Fig. 4B**). We then examined the effect of NAC, anti-ROS reagent, for its ability to suppress cell death. At 5 mM, NAC restored the cell viability up to 40% compared to less than 20% in non-treated cells (**Fig. 4C**). It also partially restored cell cycle progression and DNA synthesis (**Fig. 4D and E**). These results indicate ROS, generated in Claspin knockout cells, are also at least partially responsible for cell death.

### Claspin interacts with PI3 kinase, mTOR, and PDK1

In order to understand how Claspin may function in the PI3K/Akt/mTOR pathway, we examined whether Claspin physically interacts with the factors involved in this pathway. We expressed Flag-tagged Claspin transiently in 293T cells, and immunoprecipitated with antibody against the Flag tag. HA-tagged Claspin was used as a negative control. PI3 kinase p100β as well as mTOR were efficiently pulled down, and PI3 kinase Class III and PI3 kinase p100α were also coimmunoprecipitated (**Fig. 5A**), although the efficiency was lower. The phosphorylated form of PDK1 (S241) also interacted with Claspin. In contrast, Akt, S6 kinase and 4EBP1 did not interact with Claspin (**Fig. 5A**).

**Fig. 5.**
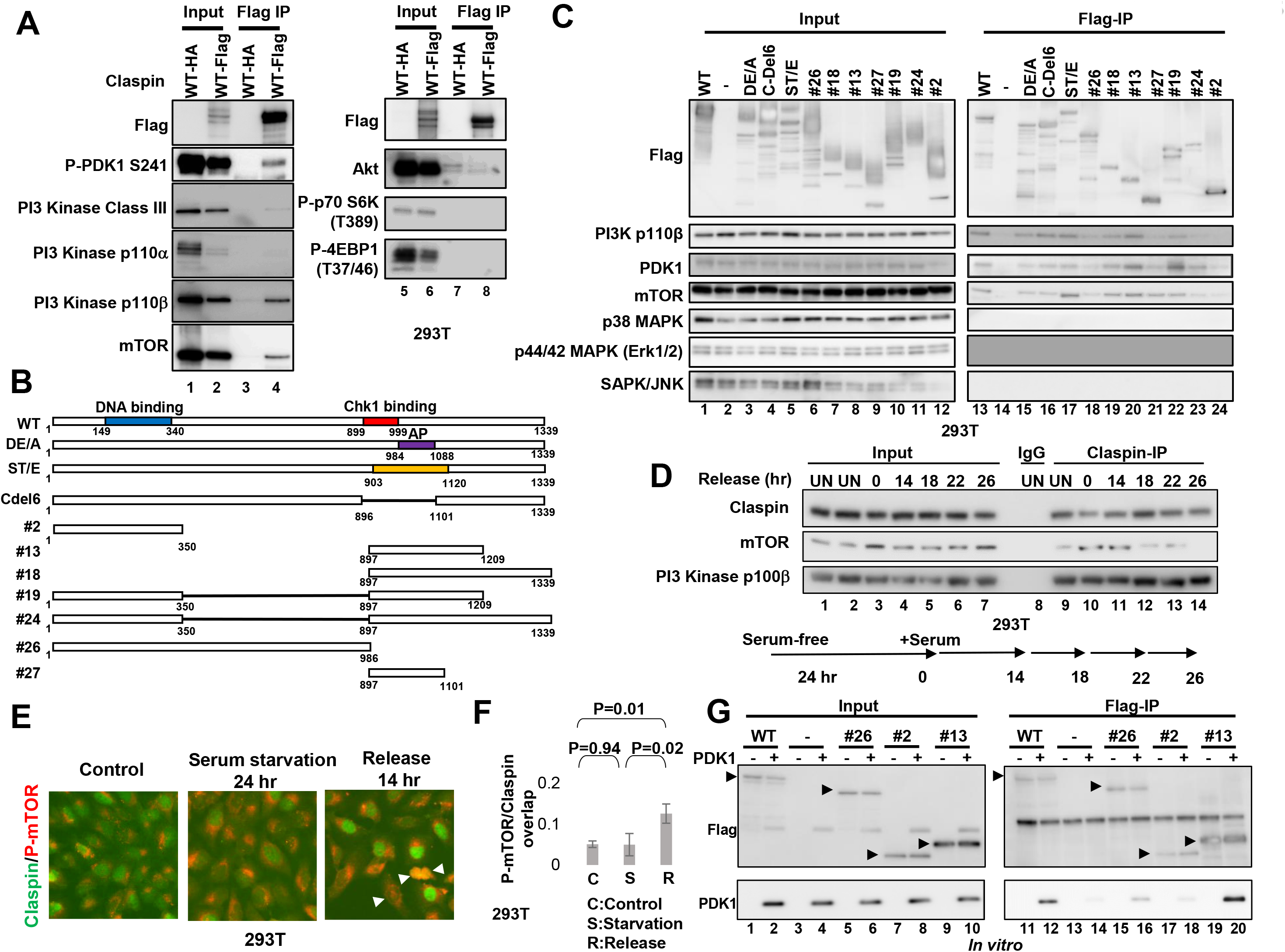
Claspin interacts with members of the PI3K-PDK1-mTOR pathway. (**A**) Flag-tagged wild-type or HA-tagged (negative control) Claspin was expressed in the 293T cells, and immunoprecipitates pulled down by anti-Flag antibody agarose beads were analyzed by western blotting with indicated antibodies. (**B**) The schematic drawing of the wild type and mutant Claspin polypeptides used in the pull-down assays. (**C**) Flag-tagged wild type or various mutant Claspin polypeptides were expressed in 293T cells, and immunoprecipitates pulled down by anti-Flag antibody agarose beads were analyzed as in (**A**). (**D**) 293T cells cultured in serum-free medium for 24 hr and released by addition of serum were harvested at indicated times. Triton-soluble extracts were prepared and were used for immunoprecipitation by anti-Claspin antibody. The input extracts and immunoprecipitates were analyzed by western blotting with antibodies indicated. UN, untreated. Lane 8, immunoprecipitated by control IgG. (**E**) 293T cells were cultured in normal medium (control) or serum-free medium for 24 hr (serum starvation 24 hr). Cells in serum-free medium were released by addition of serum for 14 hr (Release 14 hr). Cells were fixed with PFA, treated with Triton-X100, and stained with anti-mouse Alexa 488 conjugated anti-Claspin antibody (green) and anti-mouse Alexa 594 conjugated anti-P-mTOR antibody (red). Cells were observed under fluorescence microscopy. (**F**) Quantification of the fractions of P-mTOR signals overlapping with Claspin signals from (**E**). (**G**) Various Flag-tagged Claspin polypeptides were mixed with purified PDK1 *in vitro*, and pulled down by anti-Flag agarose beads, followed by analysis by western blotting with indicated antibodies.

We then next examined which domain(s) of Claspin is (are) required for interaction with these factors involved in the nutrition-induced signaling pathway. The segment containing acidic patch interacts not only with Cdc7 but also with a region of Claspin in the N-terminal region, inhibiting DNA and PCNA binding activities of Claspin (Yang et al. 2016). Various truncated segments of Claspin, C-terminally Flag-tagged, were expressed in 293T cells and immunoprecipitates were prepared by Flag antibody (**Fig. 5B**). The 897-1101aa polypeptide (#27) of Claspin could bind to mTOR and PDK1 and the 897-1209aa polypeptide (#13) could bind to PI3 kinase p110β (**Fig. 5C**, lanes 21 and 20). The N terminal polypeptides of Claspin (#2 and #26) could not bind to PI3 kinase p100β, mTOR or PDK1 (**Fig. 5C**, lanes 24 and 18). Interactions of PI3 kinase p110β, mTOR and PDK1 with #18 (897-1339aa) and #24 (1-350aa+897-1339aa) were weaker than that with #13 (897-1209aa) and #19 (1-350aa+897-1209aa), respectively (**Fig. 5C**, lanes 19/20 and 23/22), suggesting an inhibitory effect of the C-terminal 1210-1339aa segment on the interactions. The polypeptide #19 (1-350aa+897-1209aa) and #24 (1-350aa+897-1339aa) bound to PI3K p110β and mTOR less efficiently than #13 (897-1209aa) and #18 (897-1339aa), respectively, did, suggesting that the N-terminal segment may also inhibit the interaction of PI3K p110β and mTOR with the C-terminal segment of Claspin. In contrast, the N-terminal segment stimulates the interaction of PDK1 with the C-terminal segment (compare lanes 19 and 13). mTOR interacts with ST/E strongly (lane 17), suggesting that prior phosphorylation of Claspin may play a role in its interaction with mTOR.

During the recovery from serum starvation in 293T cells, Claspin interacted with mTOR. The interaction peaked at 14 hr after the release and decreased afterward. On the other hand, interaction of Claspin with PI3K was constitutively observed during the recovery from the serum starvation (**Fig. 5D**). Thiese results suggest a possibility that Claspin contributes to activation of mTOR pathway through direct protein-protein interactions.

Immunostaining indicates that mTOR is present generally in cytoplasm in asynchronously growing cells as well as in serum-starved 293T cells, whereas Claspin is present in nuclei. Upon release from the serum starvation for 14 hr, activated mTOR relocated to nuclei, overlapping with Claspin (**Fig. 5E and 5F**). Similar overlap of Claspin and activated mTOR was observed also in U2OS and HCT116 cells released from serum starvation (**Supplementary Fig. S7**). Nuclear localization of mTOR was previously reported in many cell lines including normal human fibroblasts (Zhang et al. 2002).

### PDK1 binds to phosphorylated CKBD of Claspin and facilitates binding of mTOR to Claspin

We also investigated if Claspin directly binds to PDK1 by pull-down experiments using purified Claspin and PDK1 proteins. The results indicate that Claspin directly interacts with PDK1 (**Fig. 5G**, lane 12 and **Supplementary Fig. S8A**, lane 18). A C-terminal polypeptide of Claspin (#13) bound to PDK1 directly, but N-terminal polypeptides of Claspin (#2 or #26) did only very weakly (**Fig. 5G**, lanes 16, 18 and 20). The mutant Claspin harboring ST/A or ST/E substitutions in the C-terminal segment bound to PDK1 as efficiently as the wild-type except for ST27/A which showed reduced binding to PDK1 (**Supplementary Fig. S8A**, lanes 22, 24, 26 and 28). On the other hands, the acidic domain mutant, DE/A, bound to PDK1 only weakly (**Supplementary Fig. S8A**, lane 20), largely consistent with the *in vivo* interaction (**Fig. 5C**, lane 15). Furthermore, there is a PDK1 binding site (FXXF; Devarenne TP et al., 2006) in the aa 998-1001 of the acidic domain (FGDF). These results suggest that PDK1 directly recognizes and binds to the acidic patch segment. Alternatively, PDK1 may bind preferentially to the phosphorylated forms of Claspin, since the DE/A mutant fails to recruit kinases such as Cdc7 or CK1, and is expected to be a hypo-phosphorylated form (see below).

Phosphorylated CKBD of Claspin recruits Chk1 kinase in replication checkpoint. Chk1 and PDK1 share 27.5% similarity (47.5% if confined to the N-terminal 210 amino acids) in their kinase domains. Therefore, we decided to examine the possibility that PDK1 is recruited to CKBD in phosphorylation-dependent manner. The phosphorylated CKBD polypeptide containing two critical phosphorylation sites (T916 and S945) pulled down purified PDK1 (**Fig. 6A**, lane 8). Shorter polypeptides containing only pT916 or pS945 also pulled down PDK1, albeit with reduced efficiency (**Fig. 6A**, lanes 10 and 12). With all the three peptides, prior dephosphorylation of the peptide eliminated the binding, indicating that PDK1 binds to CKBD of Claspin in a phosphorylation-dependent manner (**Fig. 6A**, lanes 9, 11,and 13). This suggests a possibility that Claspin serves as a mediator during serum-induced growth restart, and recruits PDK1 potentially as an effector kinase.

**Fig. 6.**
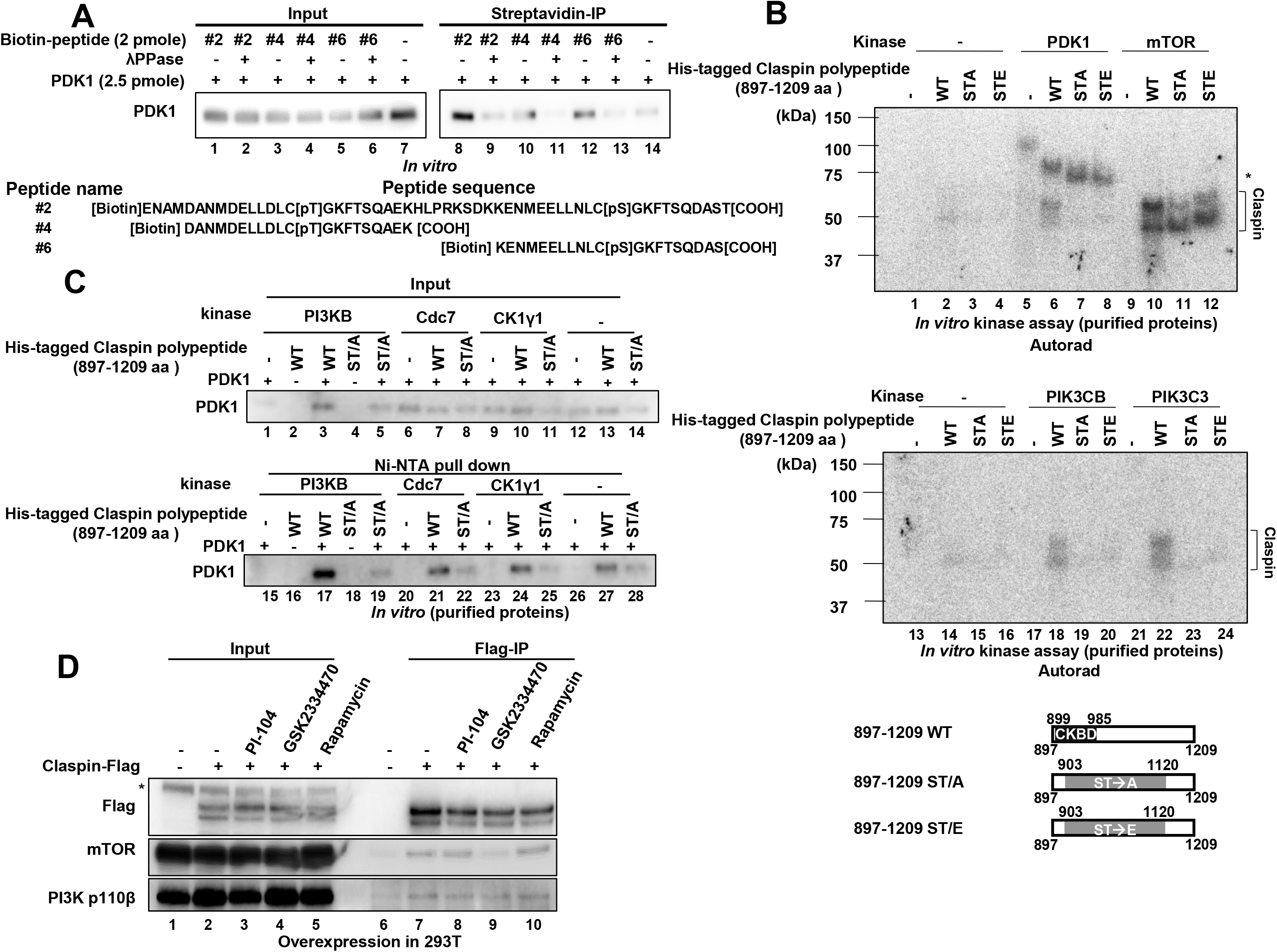
Physical and functional interactions between Claspin, PDK1, PI3K and mTOR. **(A)** Biotin-tagged CKBD peptides as indicated, treated with λPPase or not, were mixed with purified PDK1, and were pulled down by Streptavidin conjugated bead. The PDK1 co-pulled down with CKBD was analyzed by western blotting. **(B)** Purified His-tagged 897-1209aa Claspin polypeptides (wild type, ST/A and ST/E mutant) were mixed with PDK1, mTOR (upper panel) or PIK3CB, PIL3C3 (lower panel) proteins and kinase assays were conducted. Samples were analyzed on SDS-PAGE, dried and the gel was autoradiographed. CBB staining of the same gels is shown in **Supplementary Fig. S9D**. Bottom, schematic drawing of Claspin mutants used. (**C**) The 897-1209aa Claspin polypeptides (wild-type and ST/A mutant) were incubated with purified PDK1 in the presence of PI3KB, Cdc7 or CK1γ1 under the kinase reaction condition and were pulled down by Ni-NTA agarose. The co-pulled down PDK1 was analyzed by western blotting (lower panel). Upper panel, input. **(D)** Flag-tagged Claspin was overexpressed in 293T, and was treated with indicated inhibitors. The proteins pulled down by anti-Flag antibody beads were analyzed by western blotting with the antibodies indicated.

What would be the kinase responsible for phosphorylation of CKBD? We have examined the possibility that mTOR or PI3K may act as an initial kinase that stimulates PDK1 activation. Purified phosphoinositide-3-kinase beta (PIK3CB) or phosphoinositide-3-kinase, class III(PIK3C3) phosphorylated the Claspin polypeptide (897-1209aa) *in vitro*. The phosphorylation was significantly reduced by substitution of serine/ threonine with alanine in the 903-1120aa segment of Claspin (**Fig. 6B** **and Supplementary Fig. S9D**, lanes 18, 19, 22 and 23). Furthermore, PDK1, like PIK3CB and PIK3C3, also phosphorylated the 897-1209aa segment of Claspin (**Fig. 6B** **and Supplementary Fig. S9D**, lane 7). To investigate whether Claspin phosphorylated by PIK3CB promotes recruitment of PDK1 or not, we mixed his-tagged Claspin polypeptide (897-1209aa) and PDK1 in the presence of PIK3C3 or Cdc7 or CK1, the latter two of which were reported to phosphorylate CKBD of Claspin (Yang *et. al*., 2019). The Claspin polypeptide pulled down by Ni-NTA column was examined for the presence of PDK1. PDK1 interacted with the Claspin polypeptide with efficiency ∼3 times better with PIK3C3 than with Cdc7 or CK1 or without any kinase (**Fig. 6C**, lanes 17, 21, 24, and 27). Furthermore, the same polypeptide of Claspin in which serine/threonine residues in 903-1120 were replaced with alanine, interacted with PDK1 very weakly (**Fig. 6C**, lanes 18, 19, 22, 25 and 28). These results indicate that PIK3C3-mediated phosphorylation of the 903-1120 aa segment of Claspin promotes recruitment of PDK1 to Claspin. In contrast, purified mTOR phosphorylated Claspin 897-1209aa polypeptide in a manner independent of serine/ threonine residues in 903-1120aa segment (**Fig. 6B** **and Supplementary Fig. S9D**, lanes 10-12), suggesting that the 1121-1209aa segment of Claspin may be phosphorylated by mTOR.

Purified mTOR bound to the full-length Claspin, but not to the N-terminal polypeptide of Claspin (#2) (**Supplementary Fig. S8B**, lane 10), suggesting that Claspin can directly interact with mTOR through its C-terminal segment. mTOR is pulled down by the CKBD polypeptides in a phosphorylation-independent manner (data not shown). It is likely that mTOR binds to the C-terminal segments of Claspin following the recruitment of PDK1 to CKBD. This prediction is supported by the fact that the ST/E mutant of Claspin can pull down mTOR most efficiently (**Fig. 5C**, lane 17). Furthermore, binding of mTOR to Claspin was inhibited by the presence of GSK2334470, a PDK1 inhibitor (**Fig. 6D**, lane 9), suggesting that the action of PDK1 is required for mTOR binding to Claspin.

### Modification of Claspin and other factors during survival under serum starvation and following growth recovery

In both MEF and 293T cells, Claspin is mobility-shifted on SDS-PAGE after serum starvation for 24 hr and the shift disappeared after release by addition of serum (**Fig. 2A**, lane 1 **and Supplementary Fig. S9A**, lane 2). Treatment with λPPase eliminated the mobility-shift (**Supplementary Fig. S9B**, lane 4), indicating that the shift is caused by phosphorylation. Through mass-spec analyses, we have identified clusters of amino acids in the N-terminal and C-terminal segments of Claspin that are phosphorylated in response to serum starvation (**Supplementary Fig. S9C**).

Experiments with various kinase inhibitors indicated that Wortmannin (PI3K, DNA-PK/ATM inhibitor), Flavopiridol (CDKs inhibitor), GSK2334470 (PDK1 inhibitor) and XL413 (Cdc7 inhibitor) reduced the level of mobility-shift of Claspin **(Supplementary Fig. S10A**, lanes 3, 5, 8 and 9). CDK has been reported to activate mTOR to promote cell growth (Romero-Pozuelo et al., 2020). Inhibition of CDK activity by Flavopiridol did not affect the mTOR activity but reduced PDK1 activity at the time of starvation (**Supplementary Fig. S10B**, lane 5), and induced strong cell death (**Supplementary Fig. S11**). Wortmannin (PI3K, DNA-PK/ATM inhibitors) as well as Torkinib (mTOR inhibitor) caused massive cell death during growth recovery. Torkinib reduced mTOR activation and S6K/4EBP1 phosphorylation at the time of serum starvation (**Supplementary Fig. S10A**, lane 11).

GSK2334470 largely eliminated starvation-induced Claspin phosphorylation (**Supplementary Fig. S10**, lane 8), but did not induce cell death (**Supplementary Fig. S11**), and only partially inhibited mTOR and downstream kinases at the time of starvation (**Supplementary Fig. S10B**, lane 8). Cdc7 promotes DNA replication and cope with stalled forks by phosphorylating Mcm and Claspin, respectively (Yang et al., 2016, 2019). Cdc7 inhibitor, XL413, also inhibited Claspin phosphorylation (**Supplementary Fig. S10**, lane 9) but did not affect the PDK1-mTOR activity under serum starvation stress **(Supplementary Fig. S10B**, lane 9), whereas it induced cell death after serum stimulation (**Supplementary Fig. S11**). In contrast, PI3K inhibitor PI-103 did not affect Claspin phosphorylation under serum starvation but induced cell death after release from serum starvation (**Supplementary Fig. S11**), and suppressed activities of mTOR and downstream kinases at the time of starvation **(Supplementary Fig. S10B**, lane 4). The results of experiments using inhibitors support the important roles of the PI3K-mTOR pathway in serum induced growth and survival from serum starvation.

PDK1 is mobility-shifted on SDS-PAGE during serum starvation (**Supplementary Fig. S10B**, lane 2). λPPase did not affect the mobility shift (data not shown), suggesting that PDK1 undergoes some other mode of post-translational modification under serum starvation, the nature of which is currently unknown.

### Claspin promotes amino acid biosynthesis by activating the mTOR pathway in response to nutritional stress

Our results indicate that Claspin plays an important role in nutrition-induced signaling by activating the mTOR pathway to increase cell survival. In order to more precisely understand the signaling induced by serum starvation and by release from the starvation, we investigated the transcription profiles during serum starvation and after release from serum starvation by RNA-seq. Under both conditions, Claspin knockout led to significant change of transcription profiles. In contrast, we saw very little effect of Claspin knockout in 293T cells growing without any stress (**Supplementary Fig. S12**). Gene ontology analysis (GO) indicated that genes involved in translation, peptide metabolism and peptide biosynthesis are among the specifically suppressed in Claspin knockout MEF cells (**Supplementary Fig. S12**).

mTOR regulates many of the components involved in initiation and elongation stages of protein synthesis, as well as in the biogenesis of the ribosome itself (Wang & Proud; Hay & Sonenbe, 2004). mTOR also mediates translational control (Ma and Blenis, 2009) and metabolism (Sabatini, 2017). These results support the conclusion that Claspin plays crucial roles in cell growth and survival specifically during serum starvation and during growth restart from serum starvation by regulating translation and biosynthesis of ribosomes through activation of the PI3K-mTOR pathway.

## Discussion

Stress response is one of the most important biological processes in cell proliferation and survival. Although detailed studies have been conducted on cellular responses induced by various types of stresses, how these cellular responses cross talk and control cell proliferation and survival in an integrated manner has not been elucidated.

Claspin/Mrc1 is a conserved protein that mediates replication stress signal to a downstream effector kinase. Budding yeast Mrc1 has been implicated in cellular responses to osmotic, oxidative, heat and nutrient stresses in a manner independent of the replication checkpoint pathway (Duch, A. et al.2018). However, roles of Claspin in responses to similar biological stresses have been elusive in higher eukaryotes. Here, we found that Claspin plays an important role in cellular responses during the growth recovery from serum starvation.

### Claspin is required for DNA replication and survival during growth restart from serum starvation: roles in activation of PI3K-PDK1-mTOR

In Claspin knockout MEF cells, cells show reduced DNA synthesis, consistent with its role in facilitating the phosphorylation of Mcm by Cdc7 kinase. In release from serum starvation, Claspin is required for DNA synthesis, presumably reflecting its role in initiation of DNA replication. More strikingly, Claspin is required for activation of the PI3K-PDK1-mTOR pathway. The loss of the PI3K activation is not due to impaired initiation, since depletion of Cdc45, known to be essential for initiation, did not affect the viability of the cells (**Supplementary Fig. S2**). Rather, Claspin is likely to be involved in activation of the PI3K-PDK1-mTOR pathway through direct interaction with these proteins. The DE/A mutant interacted with PDK1, mTOR or PI3K only with reduced efficiency, and did not rescue the cell death in Claspin knockout cells (**Fig. 5C** **and** **Fig. 1C**). These proteins exhibited similar binding pattern to the panels of deletion derivatives of Claspin. They interact with Claspin through aa897-1209 carrying CKBD+AP segment (**Fig. 5C** **and** **Fig. 5G**).

We discovered that PDK1 binds to CKBD in a manner dependent on the phosphorylation of the critical residue in the CKBD. This is reminiscent to binding of Chk1 to the phosphorylated CKBD in response to replication stress. PDK1 may bind to phosphorylated CKBD of Claspin during recovery from serum starvation. This binding is required for further activation of mTOR, since a PDK inhibitor, GSK2334470, reduced the interaction between Claspin and mTOR (**Fig. 6D**, lane 9). Interaction of mTOR with Claspin was stimulated by the substitution of serine/ threonines in the C-terminal segment by glutamic acids (**Fig. 5C**; ST/E, lane 17). This suggests that phosphorylation of this segment of Claspin, presumably by PDK1 recruited to CKBD, facilitates the recruitment of mTOR to Claspin. On the other hand, PI3K may play a role in binding of PDK1 to Claspin, since PI3K promotes the binding of PDK1 to Claspin *in vitro* (**Fig. 6C**). However, Cdc7 or CK1γ1, which phosphorylates CKBD of Claspin, did not promote PDK1 binding to Claspin. This suggests that phosphorylation of residues other than CKBD in the 897-1209aa segment also contributes to interaction of Claspin with PDK1.

On the basis of these data, we propose that during the recovery from serum starvation, PI3K interacts with Claspin and phosphorylates the critical residues on CKBD and other segment of the 899-1120aa region. This would recruit PDK1, which then phosphorylates Claspin, facilitating the interaction of mTOR with Claspin. These results strongly suggest that Claspin may act as a platform for the activation of the PI3K-PDK1-mTOR pathway (**Fig. 7**). Currently, we do not know exactly the nature of the kinases responsible for phosphorylation of Claspin required for interaction with PDK1. Although PI3K is a prime candidate, we cannot rule out the possibility of involvement of other kinases. Cdc7 knockdown or inhibition by inhibitors induced cell death after release from starvation in 293T cells (**Supplementary Figure S2C, S11A**). This suggests a possibility that Cdc7 kinase (and CK1γ1) also contributes to phosphorylation of Claspin during growth restart.

**Fig. 7.**
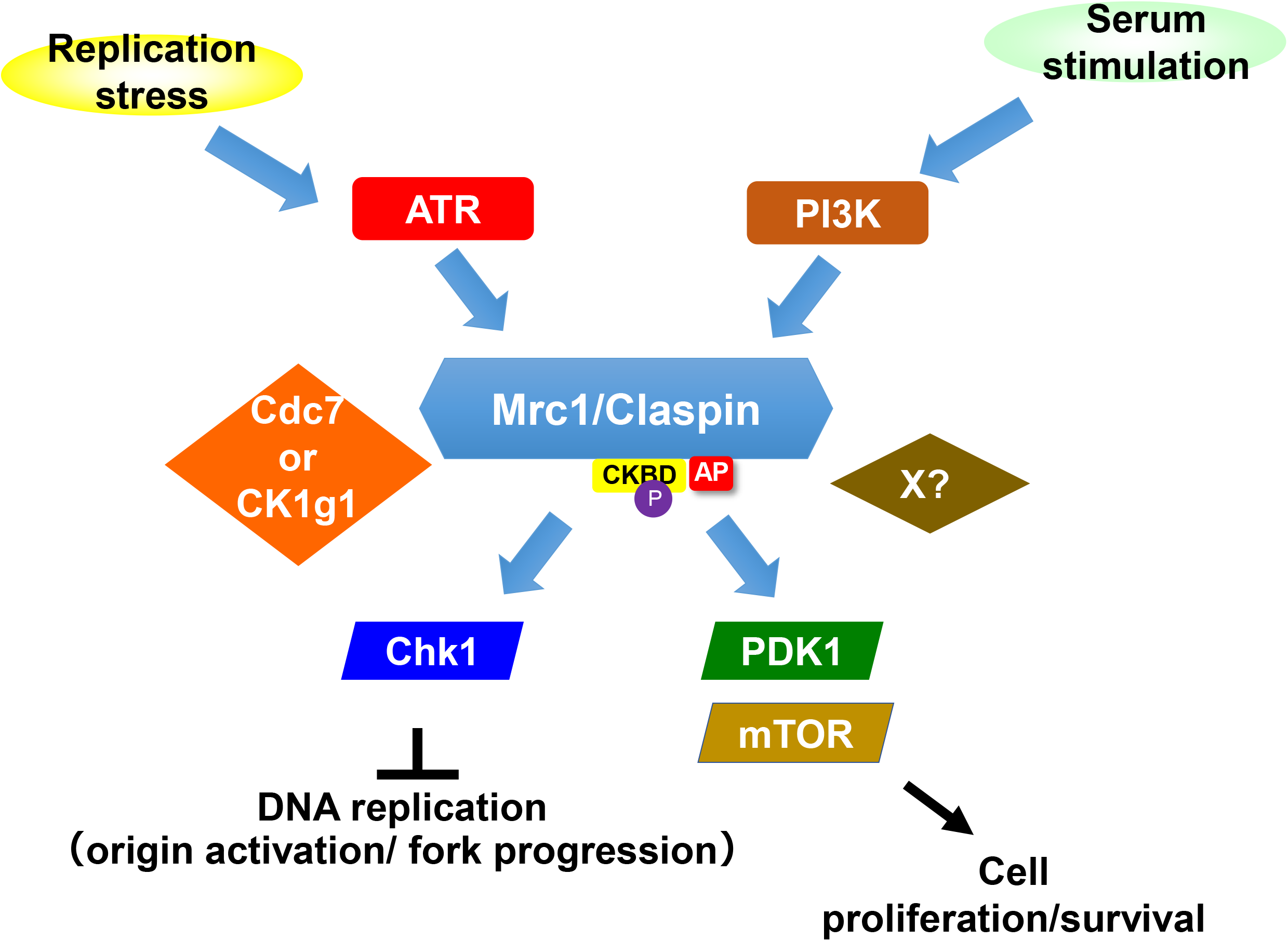
Mrc1/Claspin in serum-stimulated growth: a model. In response to serum stimulation, PI3K is activated and phosphorylates Claspin in the C-terminal segment. PDK1 binds to Claspin, and this binding is stimulated by phosphorylation with PI3K. PDK1 can bind to CKBD in a manner dependent on phosphorylation of the latter. mTOR binds to Claspin in a manner dependent on phosphorylation by PDK1. Thus, PI3K-Claspin-PDK1-mTOR pathway functions for cell growth and survival after serum stimulation. The proposed pathway is highly reminiscent to the ATR-Claspin-Chk1 pathway activated by replication stress.

### Potential activation of PI3K-PDK1-mTOR in nuclei

Activation of the PI3K pathway is a cytoplasmic event. Claspin is a nuclear protein and indeed it is detected in nuclei in 293T cells (**Fig. 5E**). How does nuclear Claspin interact with the factors involved in the PI3K-PDK1-mTOR pathway? It was reported that PI3Kβ is present also in nuclei and the nuclear PI3Kβ is essential for cell viability (Kumar et al. 2011). PI3Kβ was reported to interact with PCNA and to be required for chromatin binding of PCNA (Marqués et al. 2009). mTOR was reported to be predominantly localized in nuclei in many cell lines examined including normal human fibroblasts (Zhang et al. 2002). Thus, Claspin would activate the PI3K-PDK1-mTOR pathway in nuclei. This could contribute to the process of DNA replication and/or to the completion of mitosis.

### Loss of Claspin during growth restart from serum starvation leads to ROS and p53-dependent cell death

The transcription factor p53 is a major regulator of the DNA damage response and mTOR signaling. (A.V. Budanov, et al., 2014; P. Czarny, et al., 2015; P. Hasty, et al., 2013). The role of p53 as a responder to DNA damage may be pre-death or post-survival, depending on the extent, timing and cell type of DNA damage. Typically, severe DNA damage results in sustained activation of p53, while minor DNA damage activates p53 only briefly. Thus, the former results in cell death, while the latter induces repair and survival mechanisms (Liebl, M. C.& Hofmann, T. G, 2019). p53 is an mTOR antagonist, and thus, in response to cellular stress, p53 inhibits cell proliferation and suppresses mTOR to trigger DNA damage response. Conversely, in a negative feedback loop, mTOR increases p53 activity (K.P. Lai, et al., 2010). Furthermore, p53 also functions to deal with ROS (He & Simon, 2013). The link between ROS and p53 is complex. ROS levels are generally high in cancer cells compared to normal cells due to their hyper-metabolism, and the high ROS levels may be responsible for instability in the genome that leads to tumor formation (Achanta and Huang, 2004). Claspin depletion leads to failure to activate the PDK1-mTOR pathway, eventually leading to impaired DNA replication or unattended replication-transcription collisions. This would generate a large amount of ROS and activate p53, leading to cell death.

### Claspin promotes amino acid biosynthesis through activation of the mTOR pathway in response to nutritional stress

mTOR coordinates eukaryotic cell growth and metabolism in response to microenvironmental stimuli, including nutrients and growth factors. Extensive research during the past decades has shown that mTOR plays a crucial role also in the regulation of fundamental cellular processes such as protein synthesis, autophagy, and oxidative stress responses and that deregulated mTOR signaling is related to cancer progression and aging (Saxton et al., 2017).

By RNA-seq, we found that expression of genes involved in translation and amino acid biosynthesis decreased in Claspin knockout cells during both serum starvation and release from serum starvation (**Supplementary Fig. S12**), whereas no significant changes were observed by Claspin knockout in asynchronously growing cells. It has been reported that Mrc1 in yeasts facilitates heterochromatin formation and maintenance (Toteva et al., 2017). Potential direct roles of Claspin in regulation of gene expression are currently under investigation.

The Claspin-PI3K/PDK1/mTOR interaction is crucial for maintaining the growth from serum starvation. How does Claspin facilitate the PI3K-PDK1-mTOR pathway during growth from serum starvation? We envision two possibilities. One is that Claspin functions as an essential mediator that transmits the upstream PIKK signal to downstream effector kinases through phosphorylation relay. The other is that Claspin serves as a molecular scaffold on which the PI3K-PDK1-mTOR factors assemble and are efficiently activated. The role of such a scaffold protein has been proposed also for other signal transduction pathways or cellular reactions (Muñoz et al. 2009; Tesmer 2009).

## Conclusions

We have identified a novel role of Claspin in activation of the PI3K-PDK1-mTOR during growth restart from serum starvation. The PI3K-Claspin-PDK in serum-induced signal transmission may parallel the ATR-Claspin-Chk1 in replication stress checkpoint. PDK1 binds to phosphorylated CKBD of Claspin, like Chk1, and phosphorylates Claspin, which in turn recruits mTOR. Claspin may serve as a protein hub for activation of the PI3K-Claspin-PDK (**Fig. 7**). Our findings also point to an intriguing possibility that Claspin is a general mediator that links upstream PIKK to downstream effectors in varied environmental settings.

## Acknowledgments

We would like to thank Mayumi Shindo of Protein Analyses Laboratory of our institute for analyses of phosphorylation of the Claspin and Naoko Kakusho and Rino Fukatsu for excellent technical assistance. We also thank the members of our laboratory for useful discussion.

## Author Contributions

C-CY and HM conceived the research plans. C-CY conducted most of the experiments. HM conducted some of the plasmid constructions for expression of recombinant proteins used in this study. C-CY and HM wrote the paper.

## Materials and Methods

### Cell lines

293T, U2OS and HCT116 were obtained from ATCC. Mouse Embryonic Fibroblasts (*MEFs*) were established from E12.5 embryos as described (Yang CC et al., 2016). HCT116 were cultured at 37 °C in a 5% CO_2_ humidified incubator in McCoy’s 5A (Modified) Medium (ThermoFisher) supplemented with 10% fetal bovine serum (Gibco) and 100 U/ ml penicillin and 100 μg/ ml streptomycin. 293T, U2OS and MEF cells were cultured at 37 °C in a 5% CO_2_ humidified incubator in Dulbecco’s modified Eagle’s medium (high glucose) (Nacalai Tesque Inc.) supplemented with 15% fetal bovine serum (HANA-NESCO), 2 mM L-glutamine (Gibco), 1% sodium pyruvate (Nacalai Tesque Inc.), 100 U/ ml penicillin and 100 μg /ml streptomycin.

### Plasmid construction

The Claspin-encoding DNA fragment (*Xho*I/*Xba*I fragment) of CSII-EF MCS-mAG-TEV-His_6_-Claspin-Flag_3_, CSII-EF MCS-His_6_-Claspin-Flag_3_ or CSII-EF MCS-His_6_-Claspin-HA plasmid DNA was replaced by DNA fragments encoding portions of Claspin, amplified by PCR, to express truncated forms of Claspin (Yang CC et al., 2016). To express a Claspin mutant with an internal deletion, two PCR-amplified fragments (*XhoI*-*Bam*HI and *Bam*HI*–Xba*I or *Xho*I-*Nhe*I and *Nhe*I*– Xba*I) encoding N-terminal-proximal and C-terminal proximal segments of Claspin, respectively, were inserted at the *Xho*I-*Xba*I site of CSII-EF MCS-His_6_-Claspin-Flag_3_ to replace the Claspin insert. The *Eco*RI-*Hpa*I fragment of wild-type or mutant Claspin DNA from mAG-TEV-His-Claspin-Flag_3_ was inserted at the *Eco*RI/*Sna*BI site of pMX-IP (Addgene) to construct retroviral expression vectors. The mTOR-encoding DNA fragment amplified by PCR was inserted into CSII-EFHis_6_-MCS-Flag_3_ vector DNA (ver3.4) at the *Bam*HI site (Uno et al., 2012; Yang CC et al., 2016). Wild type and mutant 897-1209aa Claspin polypeptides were amplified by PCR and inserted into pET28a (+)TEV vector at NdeI site.

### Analyses of apoptosis

Apoptosis was analyzed by Annexin V-FITC Apoptosis Kit (Biovision). In brief, *Claspin (f/+)* and *Claspin (f/-) MEFs* were infected with Ad-*Cre* or untreated for 48 hr. Cells were passaged and cultured for further 24 hr. Cells were then placed in serum-free medium and were harvested at 0, 24 and 48 hr. Harvested cells were labeled with Propidium Iodide/RNase A solution and FITC conjugated Annexin V and incubated in the dark for 5 min at room temperature. The labeled cells were analyzed by FACS. Apoptosis was analyzed also by TUNEL assay, in which DNA fragmentation was analyzed by using In Situ Direct DNA Fragmentation Assay Kit (Abcam) according to the manufacturer’s protocol. Briefly, 1×10^5^ *Claspin(f/-) MEFs* were infected with Ad-*Cre* or untreated for 48 hr. Cells were passaged and cultured for further 24 hr. Cells were then placed in serum-free medium for 48 hr and harvested. Harvested cells were fixed with 1% paraformaldehyde and washed with PBS, followed by addition of 70% ice-cold ethanol for 30 min. After washing with wash buffer, cells were mixed with staining solution for 1 hr at 37°C and rinsed with the rinse buffer. After removing the rinse buffer by centrifugation, cells were re-suspended in Propidium Iodide/RNase A solution and incubated in the dark for 30 min at room temperature. The FITC labeled cells were analyzed by FACS.

### Senescence associated β-galactosidase activity (SA-β-gal Staining)

*Claspin (f/-) MEFs* were infected with Ad-*Cre* or untreated and cultured in serum-free medium for 48 hr. After addition of serum, cells were fixed at 0, 26 and 48 hr by 0.25% glutaraldehyde for 5 min at room temperature. After washing with PBS, cells were stained in staining solution (5 mM potassium ferrocyanide, 40 mM citrate-Na_2_HPO_4_ (pH 6.0), 2 mM MgCl_2_, 150 mM NaCl and 40 mg/mL X-gal [FUJIFILM Wako Pure Chemical Corporation]) for overnight at 37°C. After washing with PBS, cells were observed under microscopes.

### Antibodies

Antibodies used in this study are as follows. Anti-human Claspin, anti-mouse Claspin, anti-MCM4, and anti-MCM4 S6T7 were previously described (Yang CC et al., 2016). Anti-LaminB (sc-6216), anti-Chk1 (sc-8408), anti-p53 (sc-99), anti-Claspin (sc-376773) and anti-MCM2 (sc-9839) were from Santa Cruz Biotechnology; Anti-Flag (M185-3L) were from MBL; anti-Chk1 S345(#2341 or #2348), anti-Chk1 S317 (#2344), anti-mTor (#2983), anti-mTOR S2448 (#5536), anti-p44/42 MAPK (Erk1/2) (#4695), anti-SAPK/JNK (#9252), p38 MAPK (#8690), anti-p38 MAPK T180/Y182 (#4511), anti-p44/42 MAPK (Erk1/2) T202/Y204 (#4370), anti-SAPK/JNK T183/Y185 (#4668), anti-PDK1 (#3062), anti-PDK1 S241 (#3438), anti-PI3 Kinase p110β (#3011), anti-PI3 Kinase p110γ (#5405), anti-PI3 Kinase p110α (#4249), anti-PI3 Kinase Class III (#3358), anti-p70 S6 Kinase T389 (#9234), anti-4E-BP1 T37/T46 (#2855), anti-Caspase-3 (#14220), anti-Akt S473(#4058) and anti-Akt (#4691) were from Cell signaling; anti-tubulin (T5168) was from Sigma-Aldrich; anti-MCM2 S53(A300-756A) was from Bethyl; anti-p53 S37 (PAB12641) was from Abnova; anti-Histone H2A.X Ser139 (05-636) and anti-Histone H3 Ser10 (05-598) were from Merck Millipore; anti-Rb (GTX50459) was from GeneTex.

### Cell survival measurement

*Claspin (f/-) MEFs* infected with Ad-*Cre* or untreated were cultured in serum-free medium for 48 hr. After addition of serum, cells were incubated for indicated time and cell numbers were counted by staining with Trypan blue.

### Knockdown of gene expression by siRNA

siRNA used are as follows. Claspin(targeting the non-coding mRNA segment) (Sense: uuggccacugauuucaauutt; Anti-sense: aauugaaaucaguggccaatt), Cdc7 (Sense: gcagucaaagacuguggautt; Anti-sense: auccacagucuuugacugctt), CK1γ1 (Sense: ggcaauaagaaagagcaugtt; Anti-sense: caugcucuuucuuauugcctt), Tim (Sense: gagcuaagaagccuaggggtt; Anti-sense: ccccuaggcuucuuagcuctt), TopBP1 (Sense: caguggagguggaguucgutt; Anti-sense: acgaacuccaccuccacugtt), AND1 (Sense: gguguagguaacaggacauat; Anti-sense: auguccuguuaccuacaccaa), Cdc45 (Sense: gaacuugaaacugcauuuctt; Anti-sense: gaaaugcaguuucaaguuctt). MISSION® siRNA Universal Negative Control #1 (Merck) was used as a negative control. siRNAs were transfected into indicated mammalian cells using X-tremeGENE™ siRNA Transfection Reagent (Merck) for 48 hr.

### Western blotting

Protein samples were run on 5∼20% precast gradient SDS-PAGE (ATTO) or 4∼20% precast gradient gel (Shanghai WSHT Biotechnology Inc.) and then transferred to Hybond ECL membranes (GE Healthcare) followed by western blot analysis with the indicated antibodies. Detection was conducted with Lumi-Light PLUS Western Blotting Substrate (Roche) and images were obtained with LAS3000 (Fujifilm) or LAS4000 (GE).

### BrdU incorporation

To the cells in 6 wells plates, BrdU (20 mM) was added for 20 min. Cells were harvested and were fixed at -20°C by 75% ethanol. After wash with wash buffer (0.5% BSA in PBS), cells were treated with 2N HCl for 20 min and then with 0.1 M sodium Borate (pH8.5) for 2 min, both at room temperature. Then, cells were treated with FITC-conjugated anti-BrdU antibody (BD biosciences, 51-33284X or Biolegend, 364103) for 20 min at room temperature in the dark, and further incubated with propidium iodide (25 µg/ml) and RNaseA (100 µg/ml) (Sigmaaldrich) for 30 min at room temperature, followed by analyses with FACS.

### Cell fractionation and immunoprecipitation

Cells were lysed in CSK buffer (10 mM PIPES-KOH [pH6.8], 100 mM potassium glutamate, 300 mM sucrose, 1 mM MgCl_2_, 1 mM EGTA, 1 mM DTT, 1 mM Na_3_VO_4_, 50 mM NaF, 0.1 mM ATP, protease inhibitor PI tablet [Roche] and 0.5 mM PMSF, 0.1% TritonX-100 and 10 units ⁄ml Benzonase [Merckmillipore]). After rotating for 60 min at 4°C, the supernatants were recovered and were incubated with anti-Flag M2 affinity beads (SIGMA) or anti-Claspin conjugated with Dynabeads™ Protein G (Thermofisher) for 60 min at 4°C. The beads were washed with CSK buffer 3 times and proteins bound to the beads were analyzed by western blotting.

### Protein purification

Flag-tagged proteins were purified by transfection of expression plasmid DNA into 293T cells was conducted as previously described (Uno et al., 2012). Cells were lysed as above. The supernatants were mixed with anti-Flag M2 affinity beads (Sigma-Aldrich) and washed by Flag wash buffer (50 mM NaPi [pH7.5], 10 % glycerol, 300 mM NaCl, 0.2 mM PMSF and PI tablet). The bound proteins were eluted by Flag elution buffer (50 mM NaPi [pH7.5], 10% glycerol, 30 mM NaCl, 200 µg/ml 3xFlag peptide [SIGMA], 0.1 mM PMSF and PI tablet) (Uno et al., 2012). His-tagged 897-1209aa Claspin fragments were purified by transfection of expression plasmid DNA pET28a(+)TEV into *E.coli* BL21-CodonPlus. Cells were lysed by lysis buffer (50 mM Na_2_PO_4_ [pH 8.0], 300 mM NaCl and 10 mM imidazole) and mixed with Ni-NAT agarose (QIAGEN) for 1 hr at 4℃. After washing by wash buffer (50 mM Na_2_PO_4_ [pH 8.0], 300 mM NaCl and 20 mM imidazole, pH 8.0), proteins were eluted by elusion buffer (50 mM Na_2_PO_4_ [pH 8.0], 300 mM NaCl and 250 mM imidazole).

### Pull down assays with purified proteins *in vitro*

To examine the interaction between the purified Claspin (C-terminally Flag-tagged) and PDK1 or mTOR, the recombinant wild-type or mutant Claspin proteins were incubated with purified PDK1 (Carna Bioscience) or mTOR (purified as above) in a reaction buffer (40 mM HEPES-KOH [pH 7.6], 20 mM K-glutamate, 2.5 mM MgCl_2_, 1 mM EGTA, 0.01 % TritonX-100, 1 mM Na_3_VO_4_, 50 mM NaF, 1 mM ATP, 0.5 mM PMSF and PI tablet) at 4°C for 1 hr. Anti-Flag M2 affinity beads (for PDK1) or anti-Claspin conjugated dynabead Protein G (for mTOR) were added and beads were recovered, washed twice with the reaction buffer. Proteins attached to the beads were analyzed by SDS-PAGE and detected by western blotting.

### ROS assay

*Claspin* (*f/-*) cells were cultured in serum-free medium for 48 hr in a 48 well plate. After addition of serum, cells were cultured for indicated time and washed by Hanks’ balanced salt solution 2 times. After addition of DCFH-DA dye (ROS assay kit; Dojindo), cells were cultured for 30 min. After wash in Hanks’ balanced salt solution 2 times, cells were observed under fluorescence microscopy.

### Immunofluorescence

Cells were fixed with 4 % paraformaldehyde, followed by incubation for 20 min in PBS containing 0.2 % Triton X-100, which is the most popular detergent for improving the penetration of the antibody. After washing by PBS, cells were incubated in PBST/1 % BSA for 1 hr, followed by incubation with anti-Claspin and anti-mTOR antibodies in PBST/1 % BSA overnight at 4°C. After washing, cells were incubated with anti-mouse IgG (Alexa Fluor® 488) (Invitrogen) and anti-rabbit IgG (Alexa Fluor® 647) (Invitrogen) secondary antibodies in 1 % BSA for 1 hr at room temperature in the dark. After washing, cells were mounted and observed under fluorescence microscope.

### Mass spectrometry analyses of Claspin

293T cells (15 cm dish, 5 plates) were cultured in serum-containing or serum-free medium for 24 hr. Cells were lysed with RIPA buffer (50 mM Tris-HCl [pH8.0], 150 mM NaCl, 0.5 % w/v sodium deoxycholate, 1.0 % NP-40 and 0.1 % SDS) containing 1 mM DTT, 1 mM Na_3_VO_4_, 50 mM NaF, 0.1 mM ATP, 0.5 mM PMSF and PI tablet. The supernatants were mixed with anti-Claspin antibody conjugated to Dynabead protein G at 4°C for 1 hr. After washes with PBS containing 0.01 % Tween-20, the proteins on beads were separated on 4∼20% gradient gel and stained by CBB. Claspin protein band was excised from the gel, digested by Trypsin and phospho-threonines or phospho-serines were analyzed by mass spectroscopy.

### Pull down assays with CKBD peptides *in vitro*

To examine the interactions between the phosphorylated CKBD peptide of Claspin and proteins involved in the PI3K-PDK1-mTOR pathway, 200 ng biotin-tagged phosphorylated CKBD peptide of Claspin, treated with λPPase or untreated, were incubated with purified proteins in a reaction buffer (40 mM HEPES-KOH [pH 7.6], 20 mM K-glutamate, 2.5 mM MgCl_2_, 1 mM EGTA, 0.01% TritonX-100, 1 mM Na_3_VO_4_, 50 mM NaF, 1 mM ATP, 0.5 mM PMSF and PI tablet) at 4°C for 1 hr. Streptavidin beads were added and incubated at 4°C for 1 hr, then washed twice with the reaction buffer. Proteins attached to the beads were analyzed by SDS-PAGE and detected by western blotting.

### *In vitro* kinase assays of Claspin-derive polypeptides

DNA fragments coding wild-type, ST/A or ST/E mutant of Claspin polypeptides (897-1209aa) were cloned at NdeI site of pET28a(+)TEV vector, and proteins were purified by Ni-NTA agarose. The purified Claspin-derive polypeptides were incubated with the purified PDK1, mTOR, PIK3CB or PIK3C3 proteins in the kinase reaction buffer (40 mM Hepes-KOH [pH7.6], 5 mM MgCl_2_, 0.5 mM EGTA, 0.5 mM EDTA, 1 mM Na_3_VO_4_, 1 mM NaF, 2 mM DTT, 10 μM ATP, and 1 μCi [γ-ATP]) for 1 hr at 37°C. After addition of one-fourth volume of 5x sample buffer, the reaction mixtures were heated at 95°C for 1 min and analyzed by SDS-PAGE, followed by CBB staining and autoradiogram, to detect phosphorylation.

### Effects of kinase on Claspin interaction with PDK1 by pull-down assay

The purified Claspin polypeptides (897-1209aa) were incubated with the PDK1, PIK3CB, Cdc7 or CK1γ1 in the kinase reaction buffer (40 mM Hepes-KOH [pH7.6], 5 mM MgCl_2_, 0.5 mM EGTA, 0.5 mM EDTA, 1 mM Na_3_VO_4_, 1 mM NaF, 2 mM DTT, 10 μM ATP, and 1 μCi [γ-ATP]) for 1 hr at 37°C. Ni-NTA agarose were added to the reaction mixtures and the collected beads were washed with Flag wash buffer as described above. Samples were analyzed by western blotting with indicated antibodies.

### Effects of kinase inhibitors on cellular protein interactions by pull-down assay

Flag-tagged Claspin were overexpressed in 293T cells for 24 hr and then treated with DMSO (control), 2 µM PI-103, 5 µM GSK2334470 or 2.5 µM Rapamycin for 12 hr. Cells were lysed with CSK buffer as above. The supernatants were mixed with anti-Flag M2 affinity beads and the collected beads were washed with Flag wash buffer as described above. Samples were analyzed by western blotting with indicated antibodies.

### Effects of kinase inhibitors on cell viability after serum starvation and during release from serum starvation

Three sets of 293T cells were transfected with siControl or siClaspin for 24 hr in serum-free medium for 24 hr. One set was harvested at 24 hr after serum starvation. The remaining two sets were released into growth by addition of serum for 12 hr or 24 hr. Asynchronously growing 293T cells were also transfected with siControl or siClaspin for 24 hr. Inhibitors or DMSA (control) were added during the last 12 hr before the harvest. Inhibitors added and their concentrations were as follows: Wortmannin (FUJIFILM Wako Pure Chemical Corporation), 2 µM; PI-103 (Cayman Chemical Company), 2 µM; Flavopiridol (Selleck), 10 µM; UCN-01 (Cayman Chemical Company), 1 µM; VE-821(Sigmaaldrich), 10 µM; GSK2334470 (Selleck), 5 µM; XL413(AdooQ Bioscience), 10 µM; Torkinib (Selleck), 5 µM; Rapamycin (Medchemexpress), 2.5 µM and MIRIN (Sigmaaldrich), 10 µM. Cells were observed under microscope.

### RNA sequencing

RNA was isolated from *Claspin (f/-)* MEFs grown under three different conditions (grown asynchronously, grown in serum-free medium for 24 hr, and grown in serum-free medium for 24 hr and released into growth by addition of serum for 14 hr) by using RNA purification kit (QIAGEN). Purified RNAs were sequenced by GENEWIZ. Raw data were analyzed by CLC Genomics Workbench (Qiagen).

---

**Supplementary Fig. S1.**
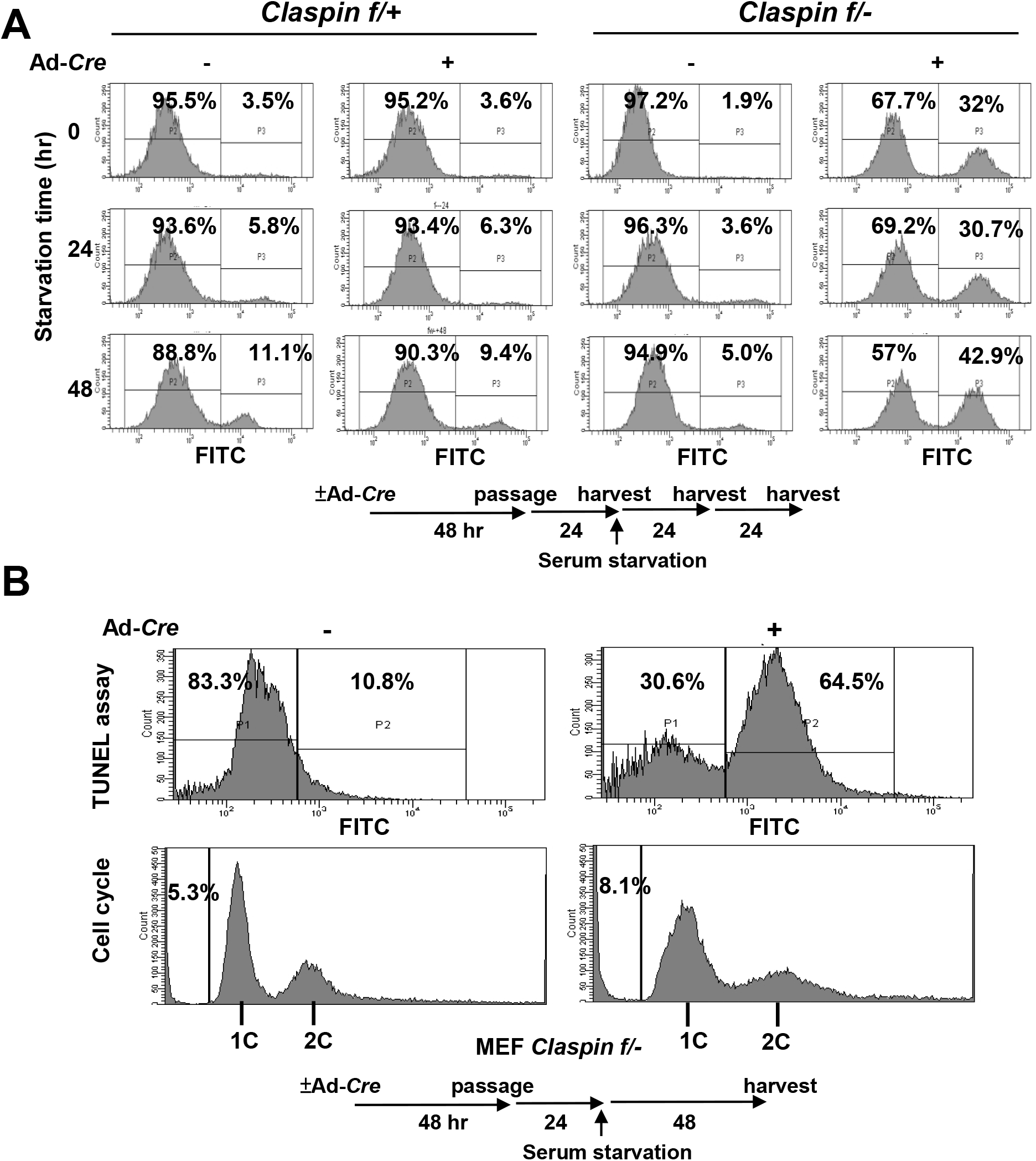
Claspin knockout MEF cells progress into apoptosis during serum starvation. **(A)** *Claspin (f/+)* and *(f/-)* MEF cells were treated with Ad-*Cre* or untreated for 48 hr. After passage and cell growth for 24 hr, cells were incubated in serum-free medium for 0, 24 and 48 hr and harvested. Cell apoptosis was analyzed by FACS after Annexin V FITC staining. P3, apoptotic cells; P2, non-apoptotic cells. (**B)** *Claspin (f/-)* MEF cells were treated with Ad-*Cre* or untreated for 48 hr. After passage and cell growth for 24 hr, cells were cultured in serum-free medium for 48 hr and harvested. Cell death was analyzed by TUNEL assay. P2, apoptotic cells; P1, non-apoptotic cells.

**Supplementary Fig. S2.**
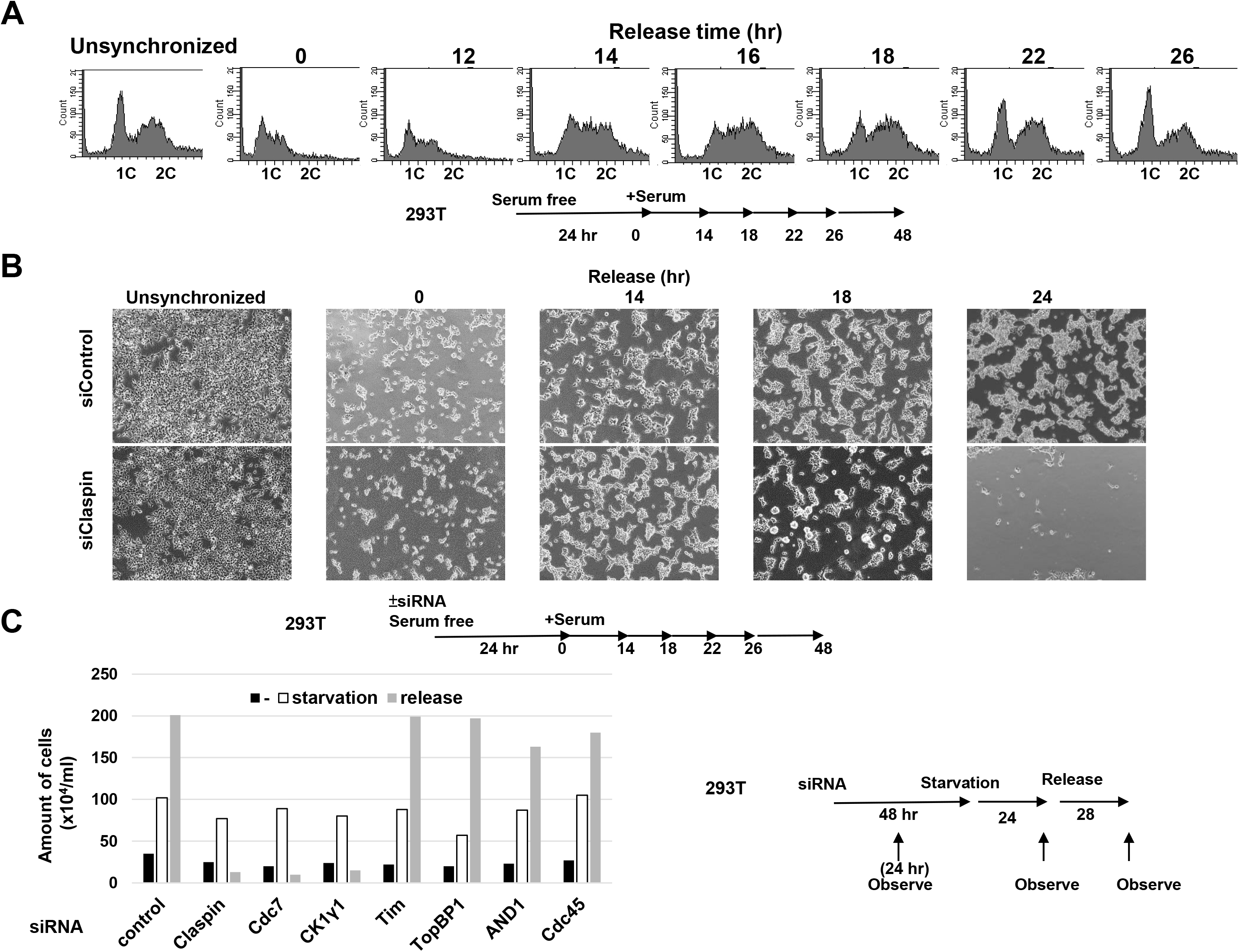
Extensive cell death is induced in Claspin knockdown 293T cells after release from serum starvation. **(A)** 293T cells cultured in serum-free medium for 24 hr and released into growth by re-addition of serum. Cells were harvested at the indicated times and cell cycle were analyzed by FACS. (**B**) siControl- or siClaspin-treated 293T cells were cultured in serum-free medium for 24 hr and released into growth by re-addition of serum. Cells were observed under microscope at the indicated times. Unsynchronized, cells treated with siControl or siClaspin for 48 hr.

**Supplementary Fig. S3.**
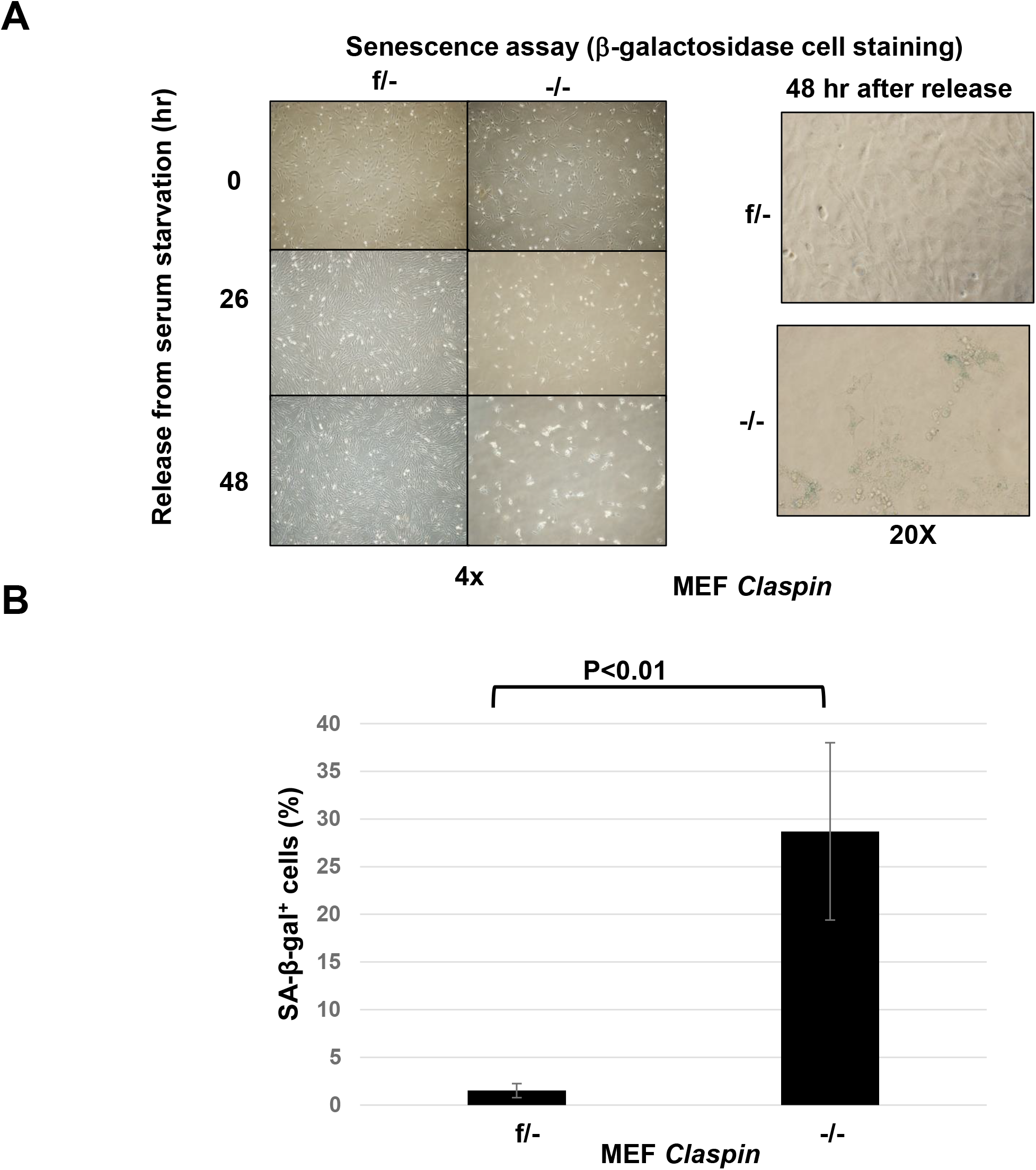
Claspin knockout MEF cells progress into senescence after release from serum starvation. **(A)** *Claspin (f/-)* MEF cells treated with Ad-*Cre* or untreated were cultured in serum-free medium for 48 hr, and were released into growth for 0, 26 or 48 hr. Photos were taken at each timepoint, and at 48 hr, cells were stained by β-galactosidase and observed under microscope. Left, 4x magnification; right, 20x magnification. (**B)** Quantification of β-galactosidase positive cells in **A**. Experiments were conducted three times and error bars are indicated.

**Supplementary Fig. S4.**
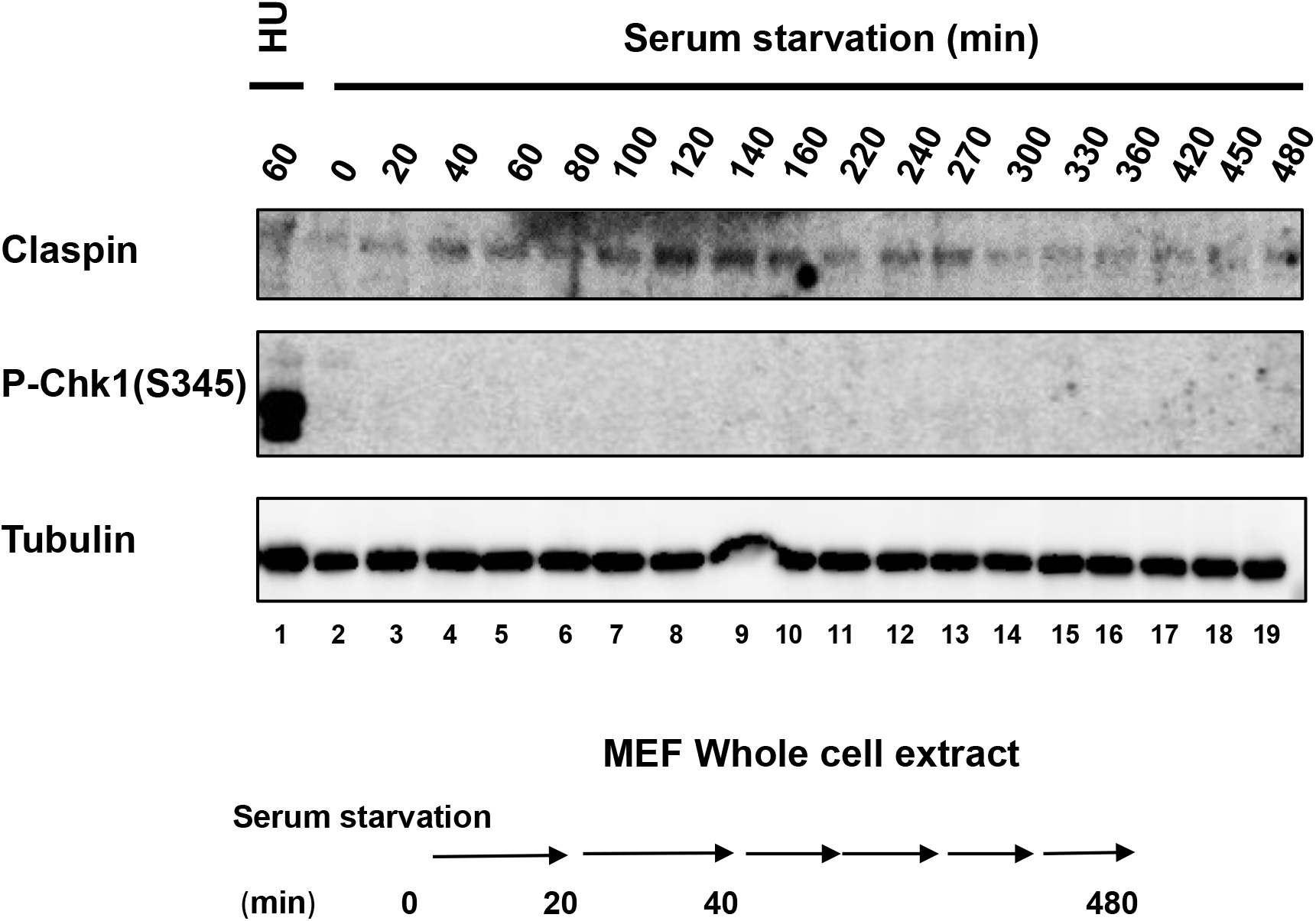
Replication checkpoint is not activated in release from serum starvation in MEF cells. MEF cells cultured in the serum-free medium and harvested at indicated times. Samples were analyzed by western blotting with the indicated antibodies. The phosphorylated Chk1 (S345) is an indicator for replication checkpoint activation. MEF cells treated with 2 mM HU for 60 min (lane 1) was used as a positive control.

**Supplementary Fig. S5.**
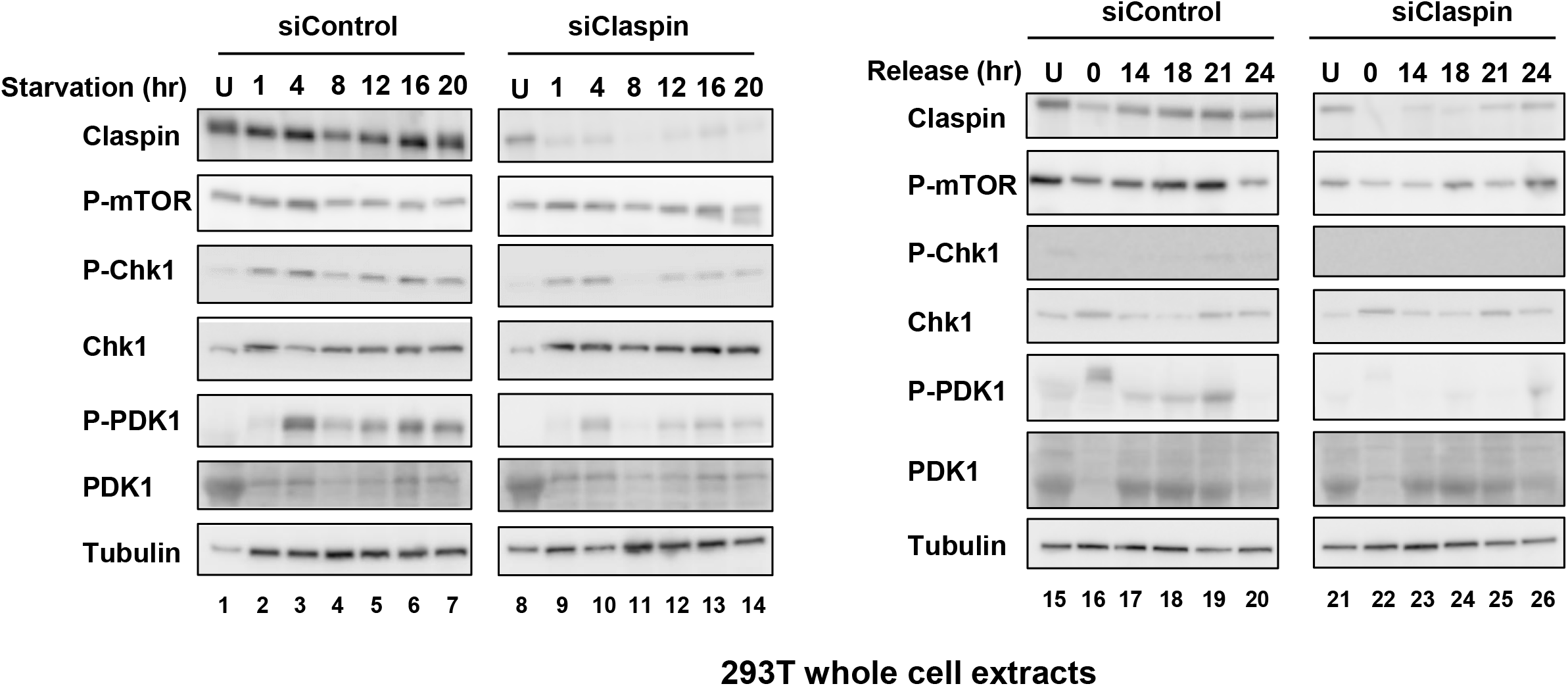
The Claspin-PDK1-mTOR pathway is important for growth recovery from serum starvation in 293T cells. **(A)** 293T cells were treated with siControl or siClaspin for 24 hr, cultured in serum-free medium and harvested at indicated times (left). After 24 hr of serum starvation, cells were released by addition of serum and harvested at indicated times (right). The whole cell extracts were analyzed by western blotting with indicated antibodies.

**Supplementary Fig. S6.**
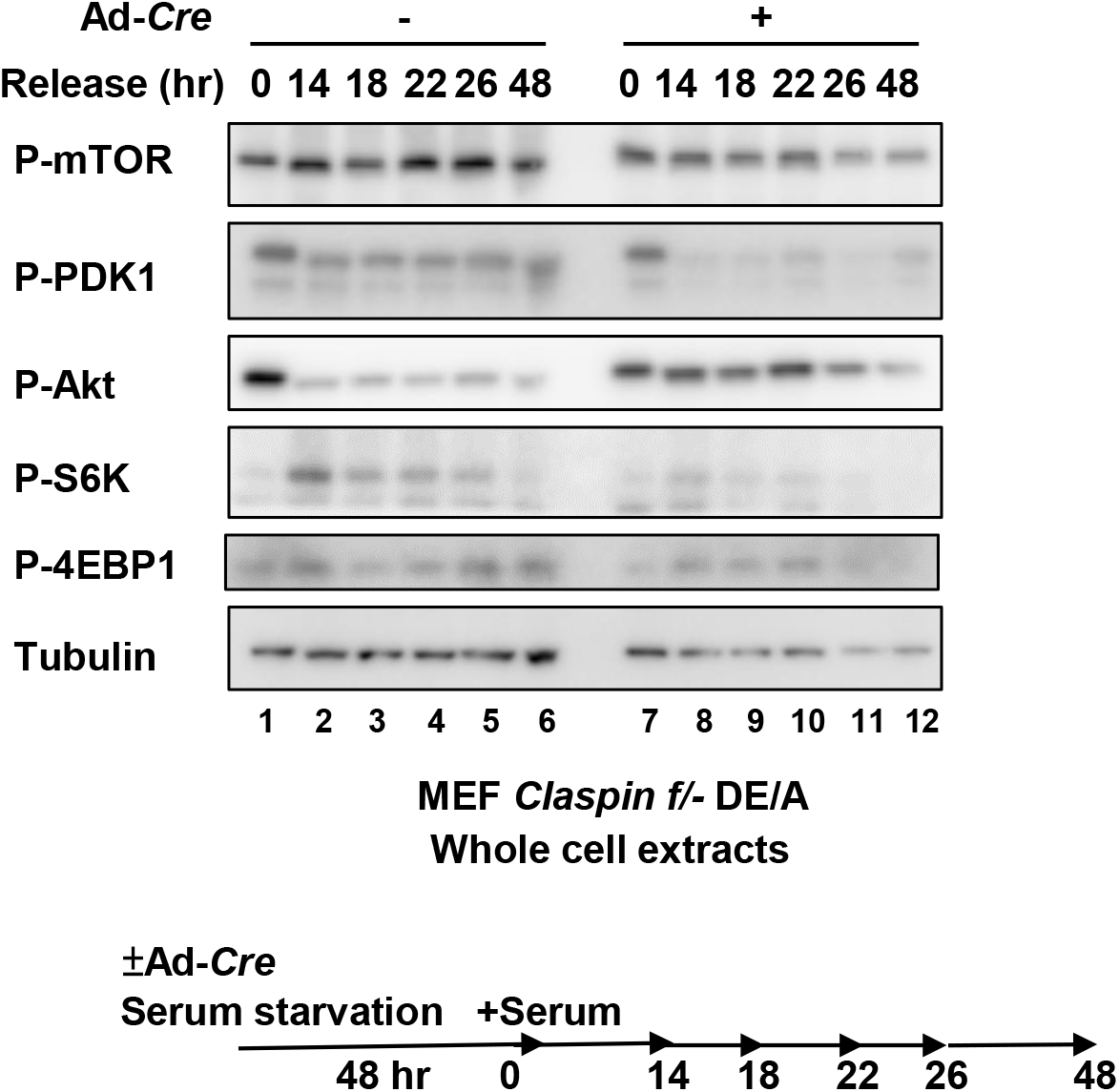
AP is required for growth recovery from serum starvation. *Claspin (f/-)* MEF cells exogenously expressing the DE/A mutant were treated with Ad-*Cre* or untreated and cultured in serum-free medium for 48 hr. Growth was restarted by addition of serum and cells were harvested at indicated times. The whole cell extracts were analyzed by western blotting with indicated antibodies.

**Supplementary Fig. S7.**
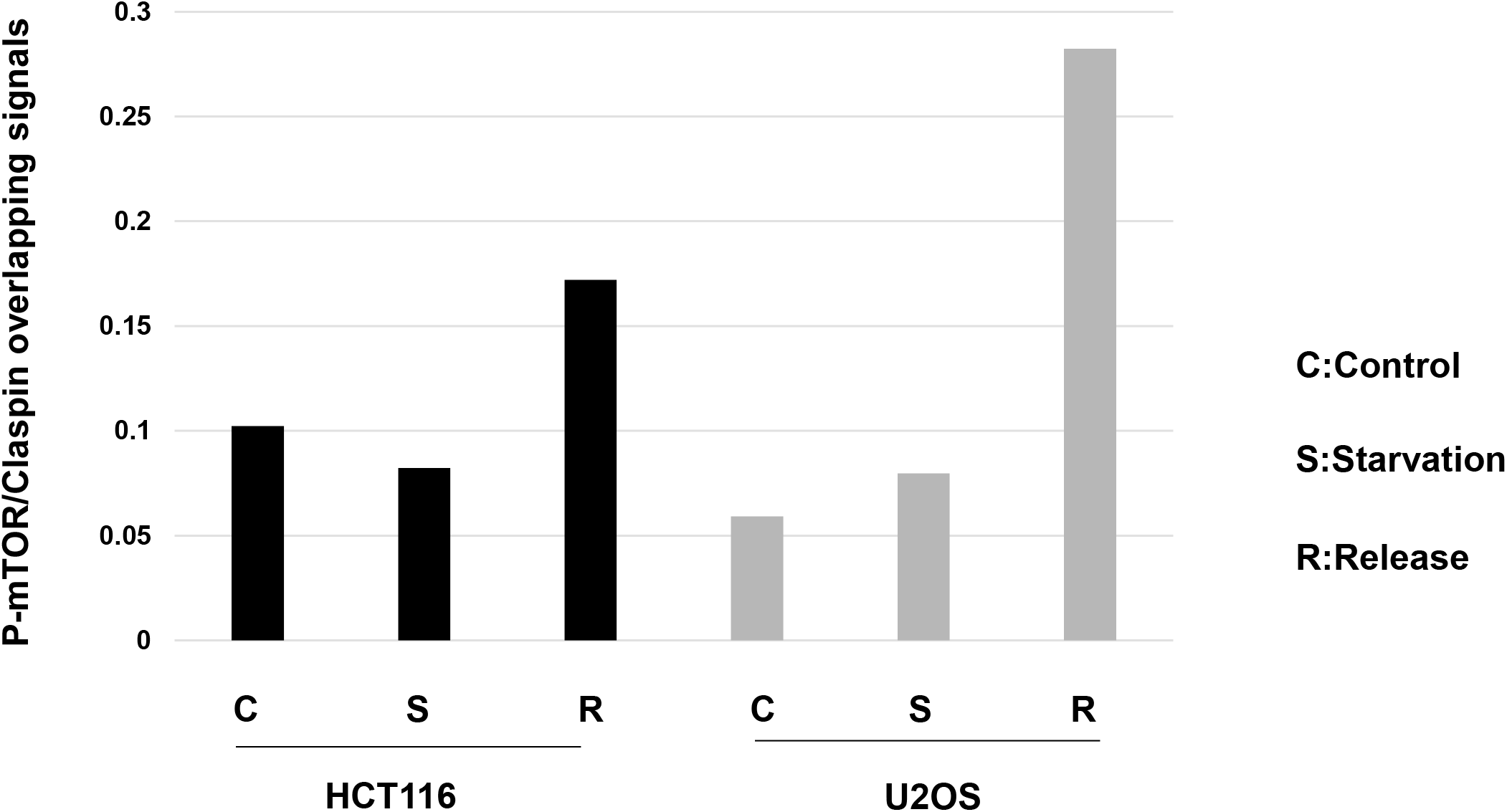
Nuclear colocalization of mTOR and Claspin in cells released from serum starvation. HCT116 and U2OS cells were cultured in normal medium or serum-free medium for 24 hr. A portion of cells in serum-free medium was released by addition of serum for 14 hr. After treating the fixed cells with Triton, Alexa 488 conjugated with anti-Claspin antibody and anti-mouse Alexa 594 conjugated with anti-P-mTOR (phosphorylated at S2448) were used to detect the signals. Cells were observed under fluorescence microscopy. The fractions of P-mTOR signals that overlap with the Claspin signals were quantified in cells under normal growth, starvation, or released from starvation.

**Supplementary Fig. S8.**
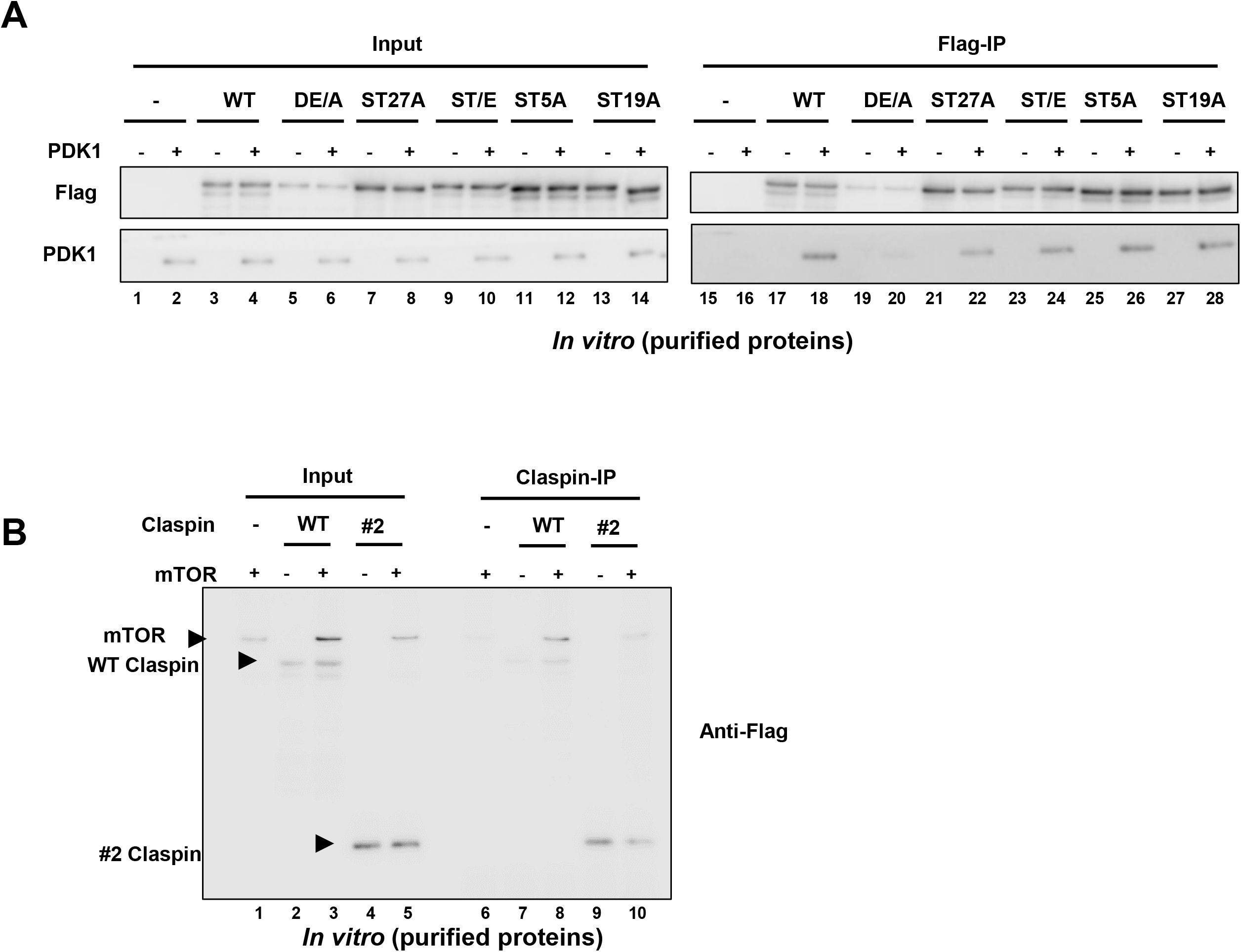
Interaction of Claspin with PDK1 and mTOR *in vitro*. Purified PDK1 (**A**) or mTOR (**B**) was incubated with various purified Flag-tagged Claspin polypeptides, pulled down by anti-Flag agarose beads and was analyzed by western blotting with indicated antibodies. One fifth of the reaction mix before pull-down is analyzed as Input.

**Supplementary Fig. S9.**
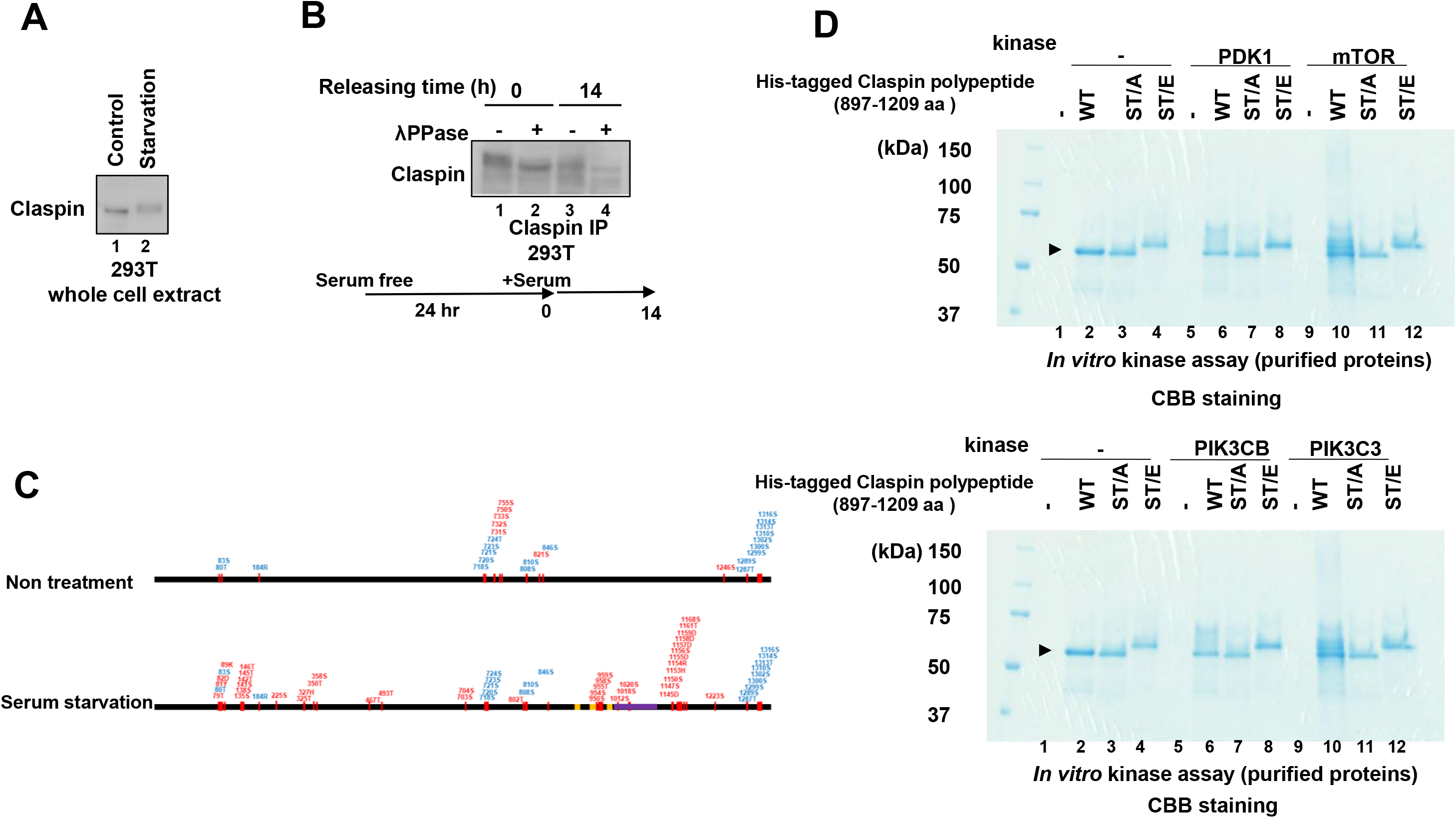
Serum starvation induces Claspin phosphorylation by PI3K pathway. (**A**) 293T cells were starved for serum for 24 hr. The whole cell extracts were analyzed by western blotting using anti-Claspin antibody. (B) 293T cells were serum-starved for 24 hr, released for 14 hr and harvested. Triton-soluble extracts were treated with λPPase for 30 min (+) or untreated (-), and then analyzed by western blotting with anti-Claspin antibody. (**C**) Phosphorylation sites of Claspin in the non-treated cells and in serum-starved (24 hr) cells were determined by mass spectrometry analyses and are shown on the drawing along the polypeptide map. Phosphorylation sites identified are indicated by small bars and numbers. Red and blue numbers indicate those unique to each condition and those common to both conditions, respectively. Three Chk1 binding domains (CKBD) and AP (acidic patch) are shown by yellow boxes and by a purple box, respectively. (**D**) CBB staining pattern of the kinase assays shown in Fig. 6B. The wild-type and mutant Claspin polypeptides (897-1209 aa) were incubated with PDK1 or mTOR (upper panel) or PIK3CB or PIK3C3 (lower panel) proteins and kinase assays were conducted. Samples were analyzed on SDS-PAGE and stained by CBB.

**Supplementary Fig. S10.**
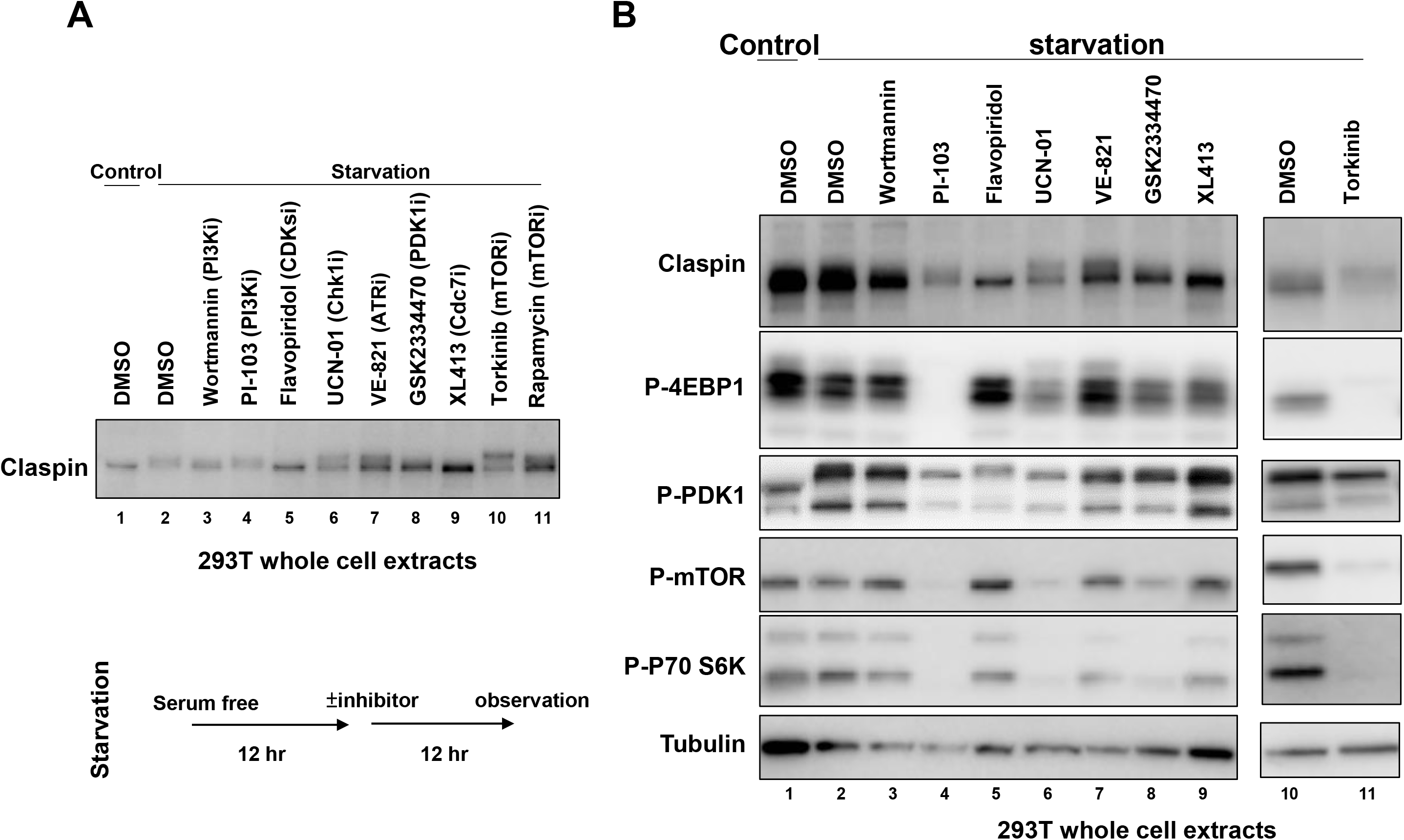
Effects of inhibitors on various proteins after serum starvation. (**A**) 293T cells, serum starved for 12 hr, were treated with inhibitors for another 12 hr under the same condition. Whole cell extracts were analyzed by western blotting with anti-Claspin antibodies. (**B**) 293T cells were starved for serum for 24 hr, and inhibitors shown were added during the last 12 hr before the harvest. The whole cell extracts were analyzed by western with the antibodies indicated. DMSO was added as a control.

**Supplementary Fig. S11.**
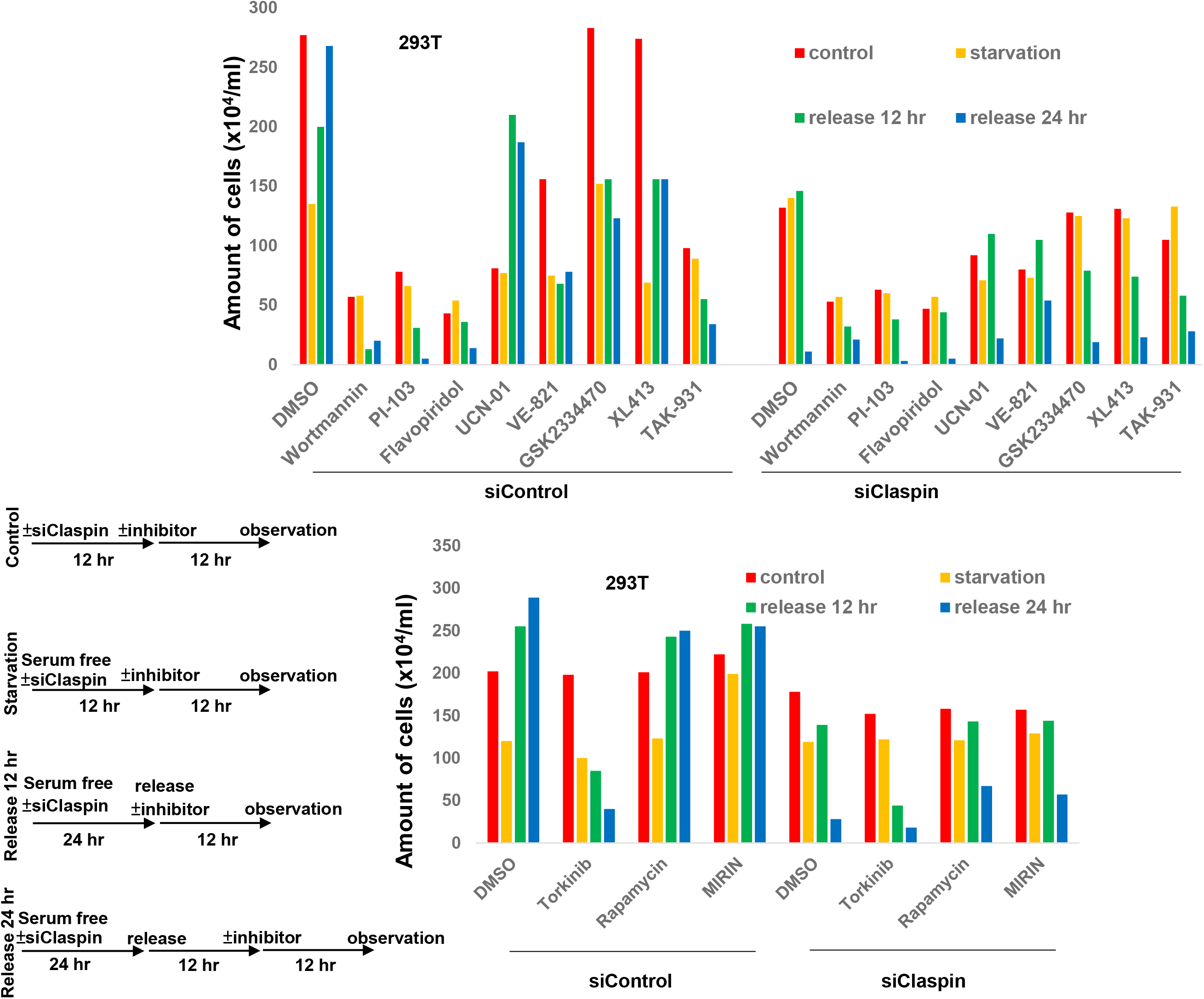
Effects of kinase inhibitors on growth of 293T cells starved for serum and on that of those released from serum starvation. 293T cells were treated with siControl or siClaspin for 24 hr in normal medium (Control), or in serum-free medium (Starvation), or released for 12 hr (Release 12 hr) or 24 hr (Release 24 hr) after 24 hr serum starvation. Inhibitors, as indicated, or DMSO (control) were added for the final 12 hr before the harvest. Cells were observed under microscope and live cell numbers were counted.

**Supplementary Fig. S12.**
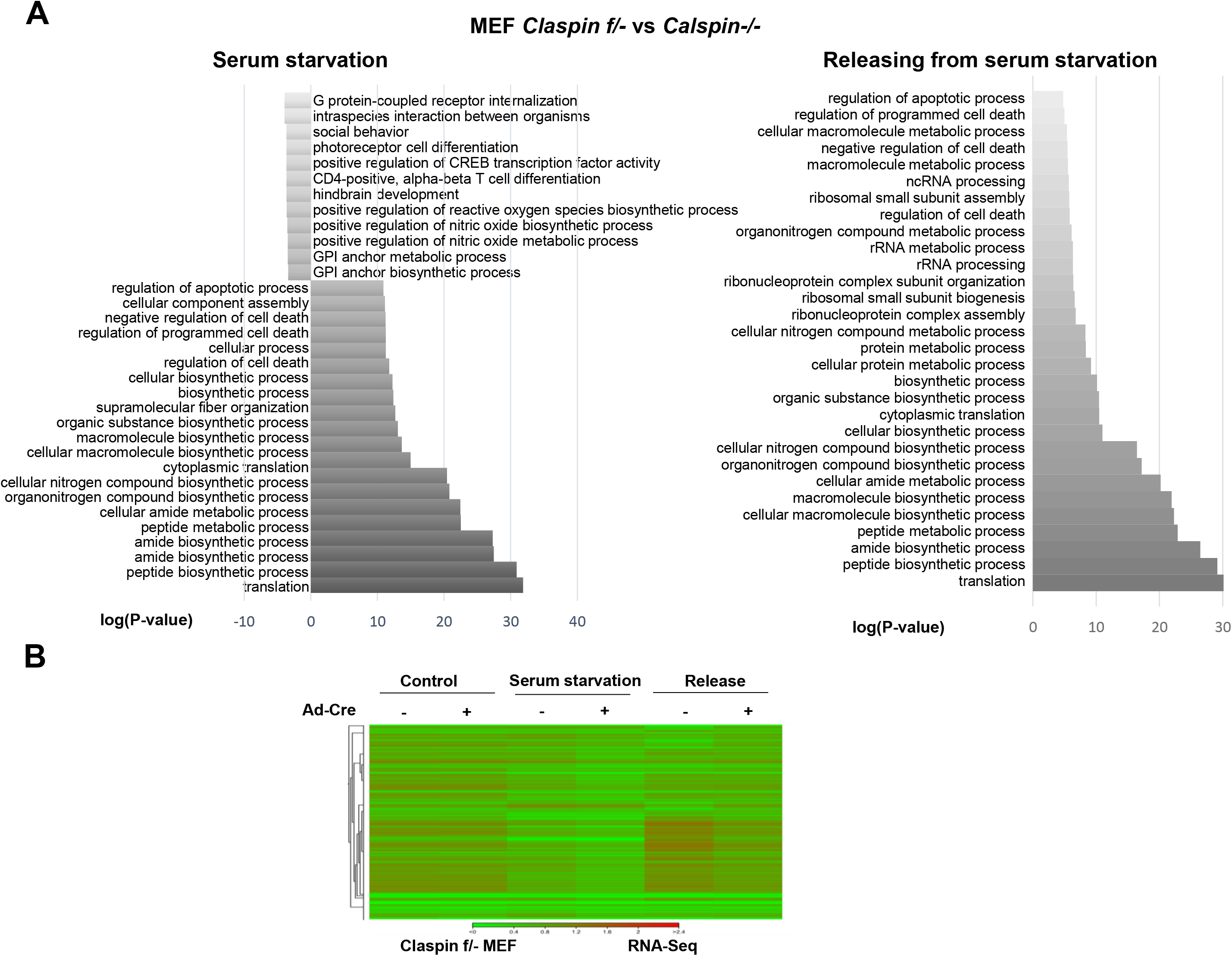
Effects of Claspin knockout on transcription profiles during serum starvation and during the release from starvation. RNA was isolated from *Claspin (f/-)* MEF, treated with Ad-*Cre* or untreated, grown under three different growth conditions (grown asynchronously, grown in serum-free medium for 24 hr, and released from serum starvation for 14 hr). **(A)** Enriched GO functions of differentially expressed genes are presented. log (p-value)>0, genes upregulated in *Claspin(f/-)*; log(p-value) <0, genes downregulated in *Claspin (f/-)*. GO, gene ontology. (**B**) Heat map illustrating differential expression data obtained by RNA-Seq. Pairwise comparisons are shown for each apricot cultivar (columns). Red, positive log fold-change (log FC) indicating higher expression in the untreated cells compared with Ad-*Cre* treated cells; green, negative log FC.

