## Supplemental material for "Claspin is required for growth recovery from serum starvation through regulating the PI3K-PDK1-mTOR pathway"

Running title:
Claspin in growth from quiescence

*Correspondence should be addressed to
Hisao Masai


### Materials and Methods

#### Cell lines

*Cre* treated cells; green, negative log FC.

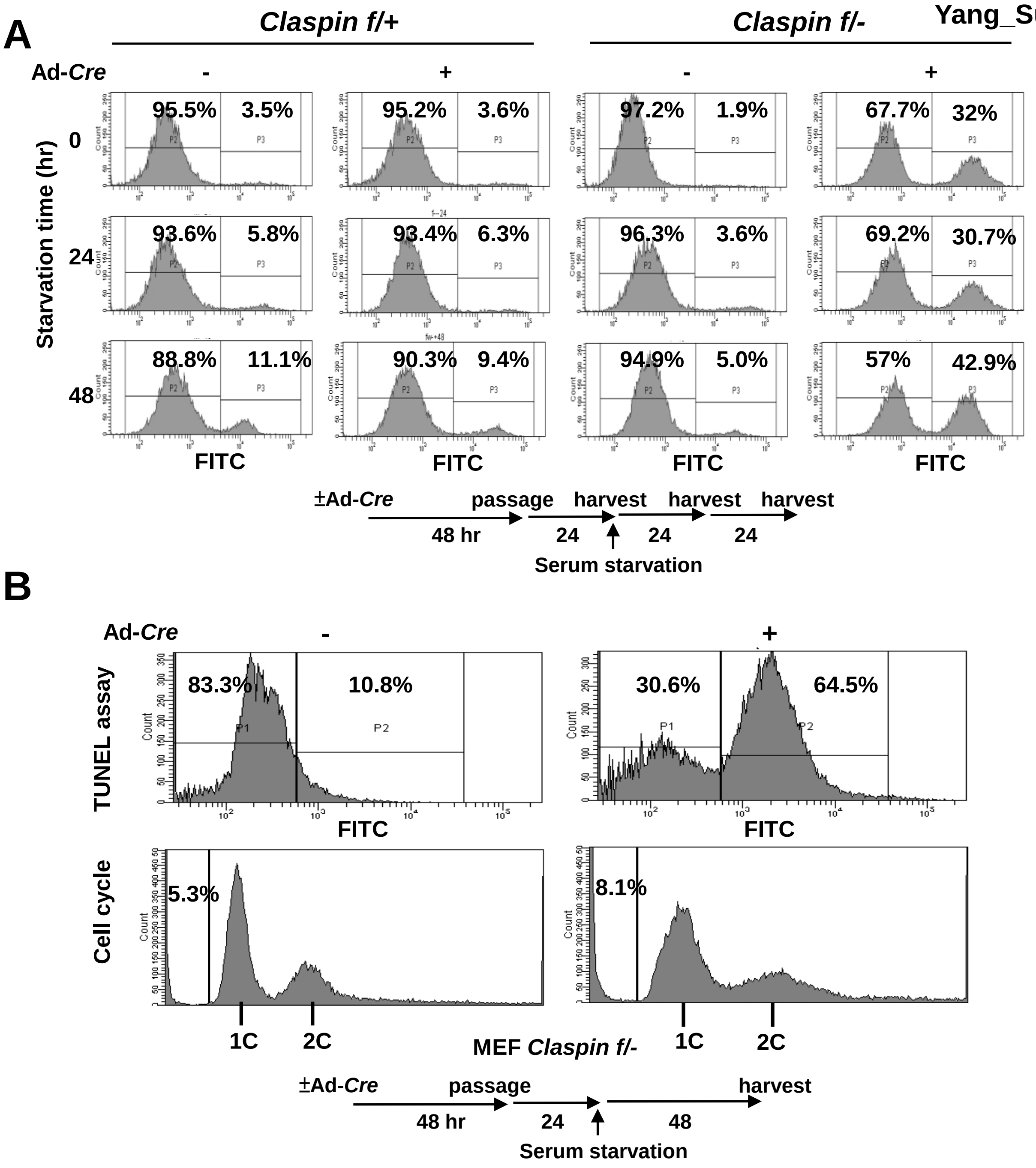
A
Claspin f/+
Claspin f/-
Ad-Cre
-
+
-
+
Starvation time (hr)
0
24
48
95.5%
3.5%
95.2%
3.6%
97.2%
1.9%
67.7%
32%
93.6%
5.8%
93.4%
6.3%
96.3%
3.6%
69.2%
30.7%
88.8%
11.1%
90.3%
9.4%
94.9%
5.0%
57%
42.9%
FITC
±Ad-Cre
passage
harvest
harvest
harvest
48 hr
24
24
24
Serum starvation
B
Ad-Cre
-
+
TUNEL assay
83.3%
10.8%
30.6%
64.5%
FITC
Cell cycle
5.3%
8.1%
1C
2C
MEF Claspin f/-
±Ad-Cre
passage
harvest
48 hr
24
48
Serum starvation

Yang_Supplementary Figure S2

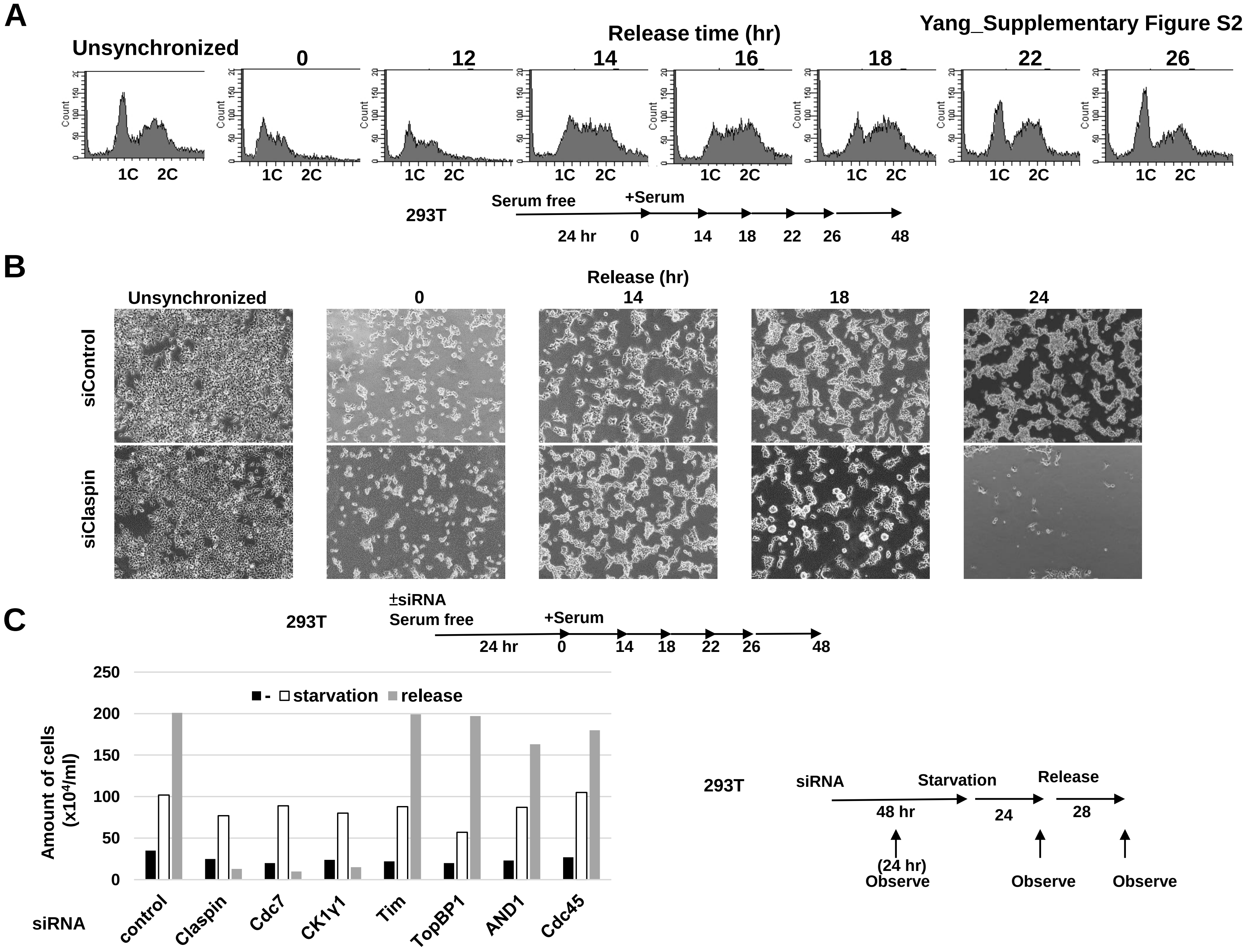

A

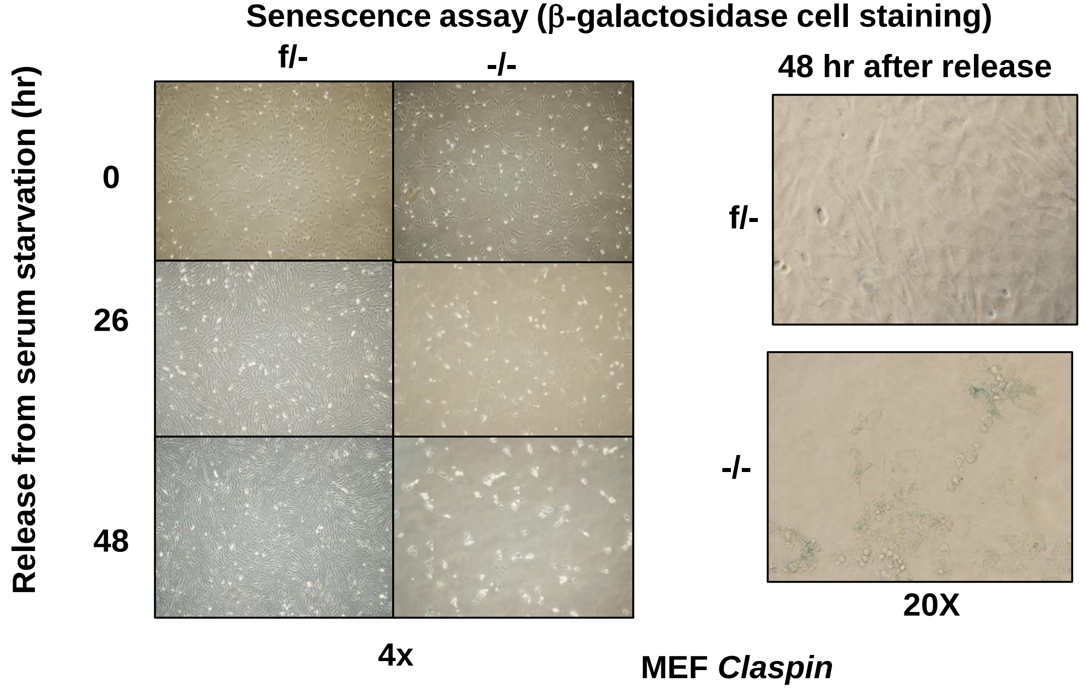
Senescence assay (β-galactosidase cell staining)
f/-
-/-
Release from serum starvation (hr)
0
26
48
4x
48 hr after release
f/-
-/-
20X
MEF Claspin

B

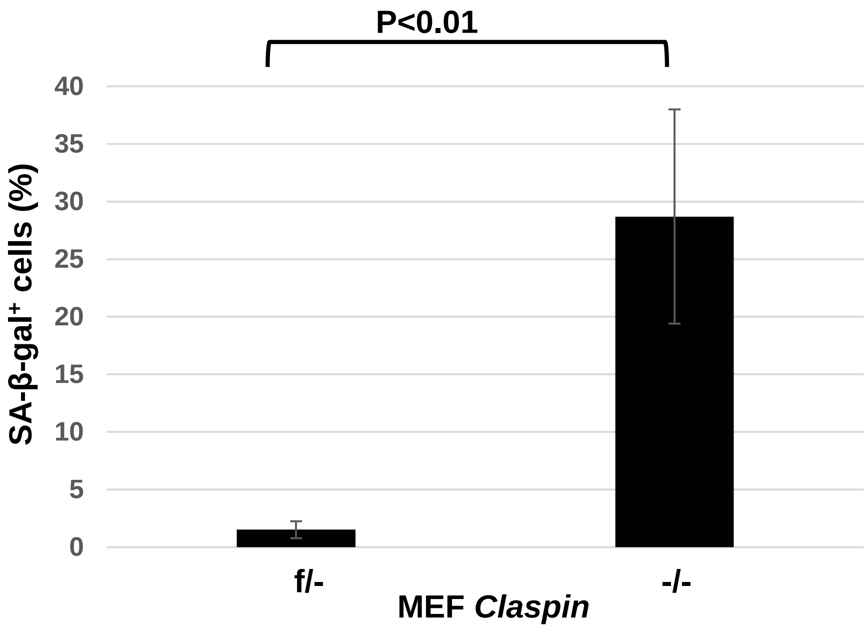
P<0.01
SA-β-gal⁺ cells (%)
40
35
30
25
20
15
10
5
0
f/-
-/-
MEF Claspin

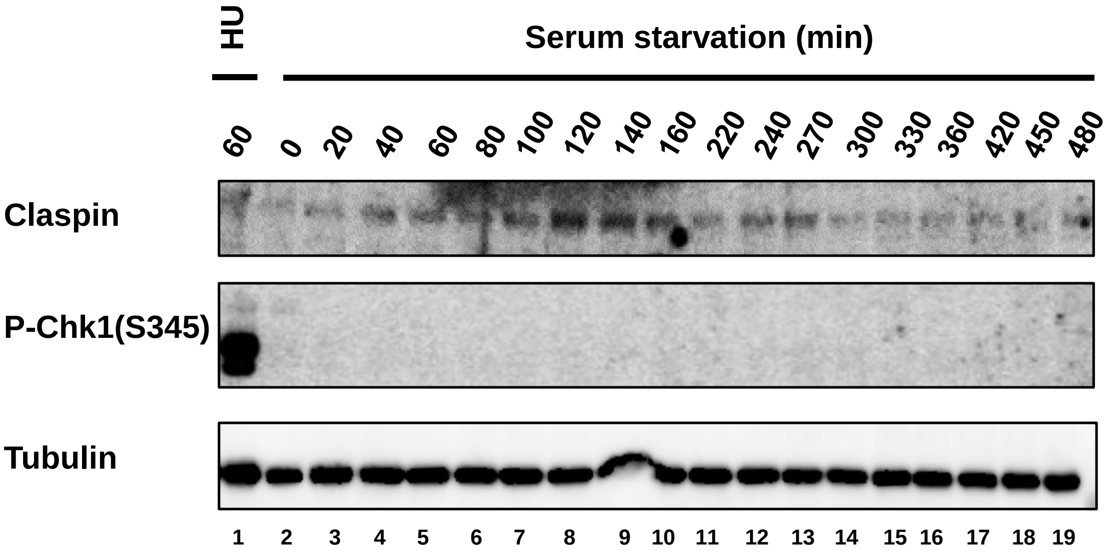


MEF Whole cell extract

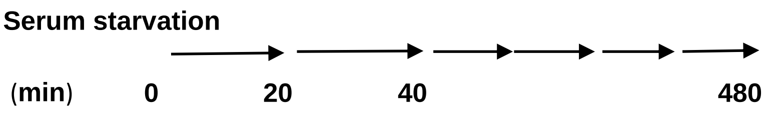

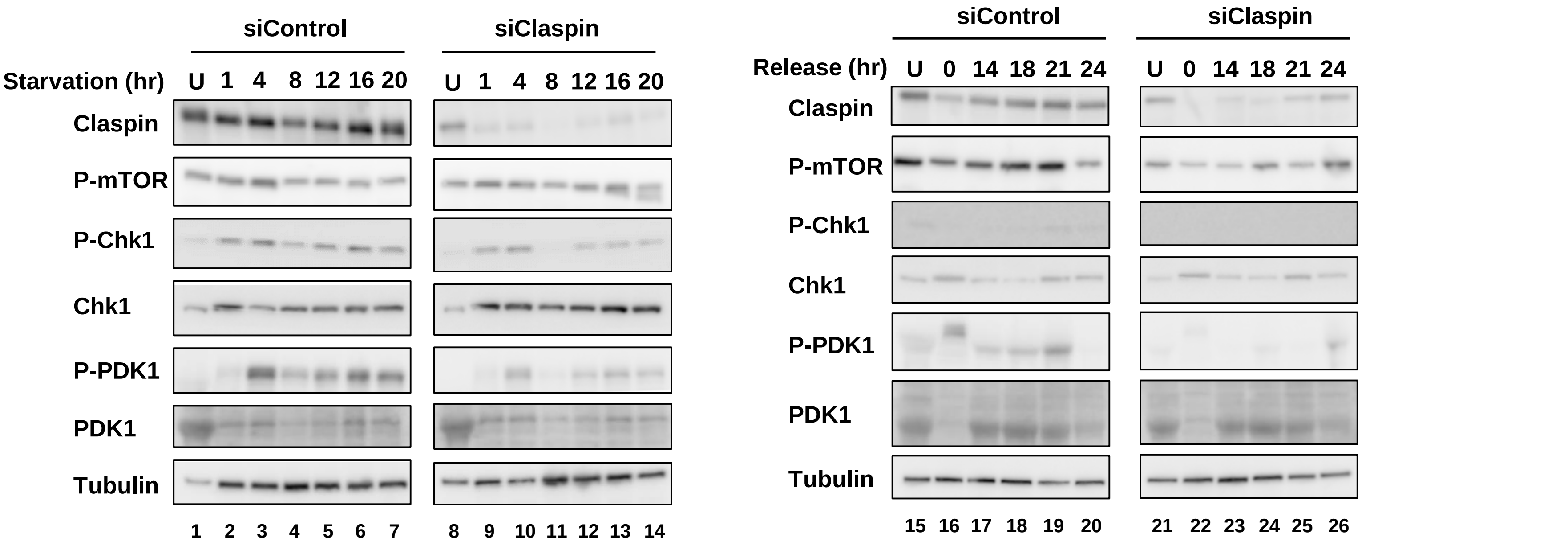


293T whole cell extracts

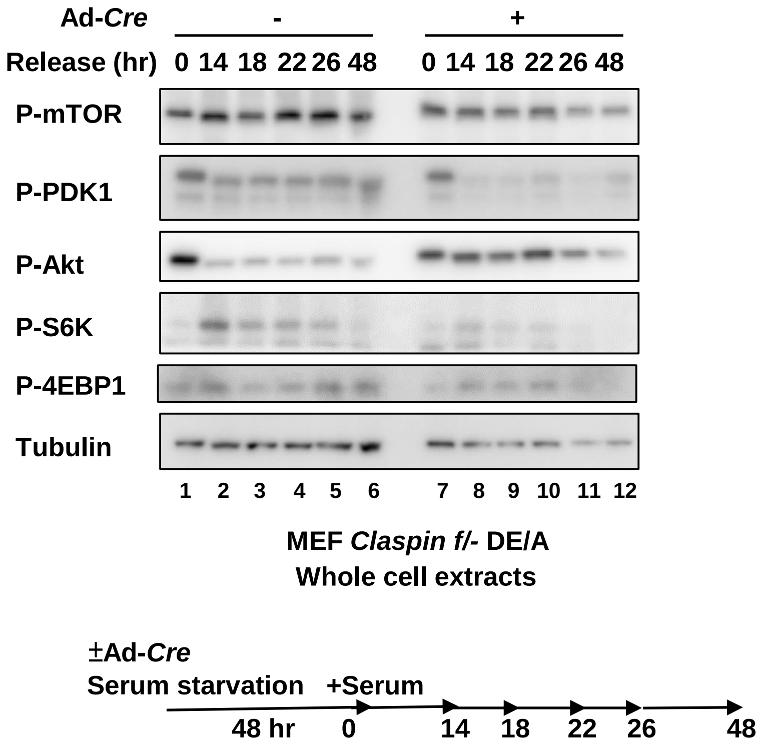

Ad-*Cre*
-
+
Release (hr)
0 14 18 22 26 48
0 14 18 22 26 48
P-mTOR
P-PDK1
P-Akt
P-S6K
P-4EBP1
Tubulin
1 2 3 4 5 6 7 8 9 10 11 12
MEF *Claspin f/-* DE/A
Whole cell extracts
±Ad-*Cre*
Serum starvation
+Serum
48 hr
0
14
18
22
26
48

Yang_Supplementary Figure S7

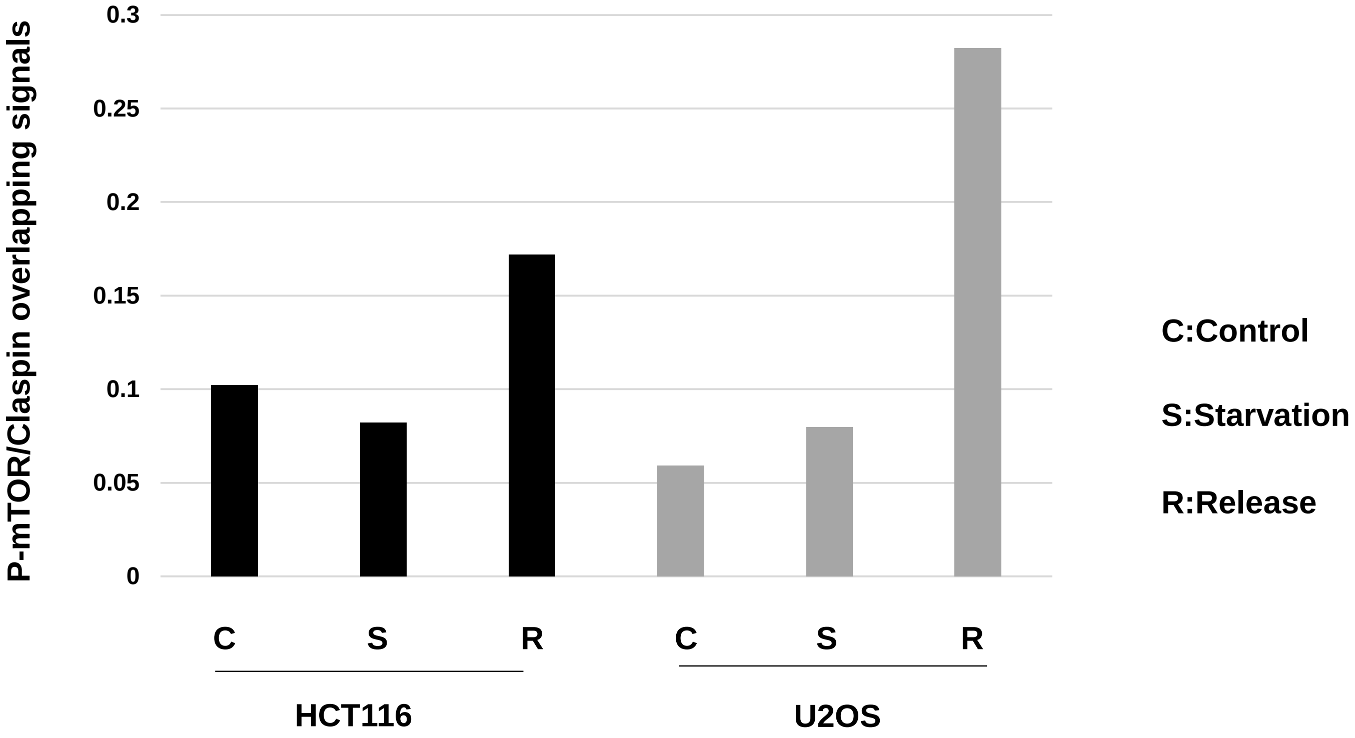

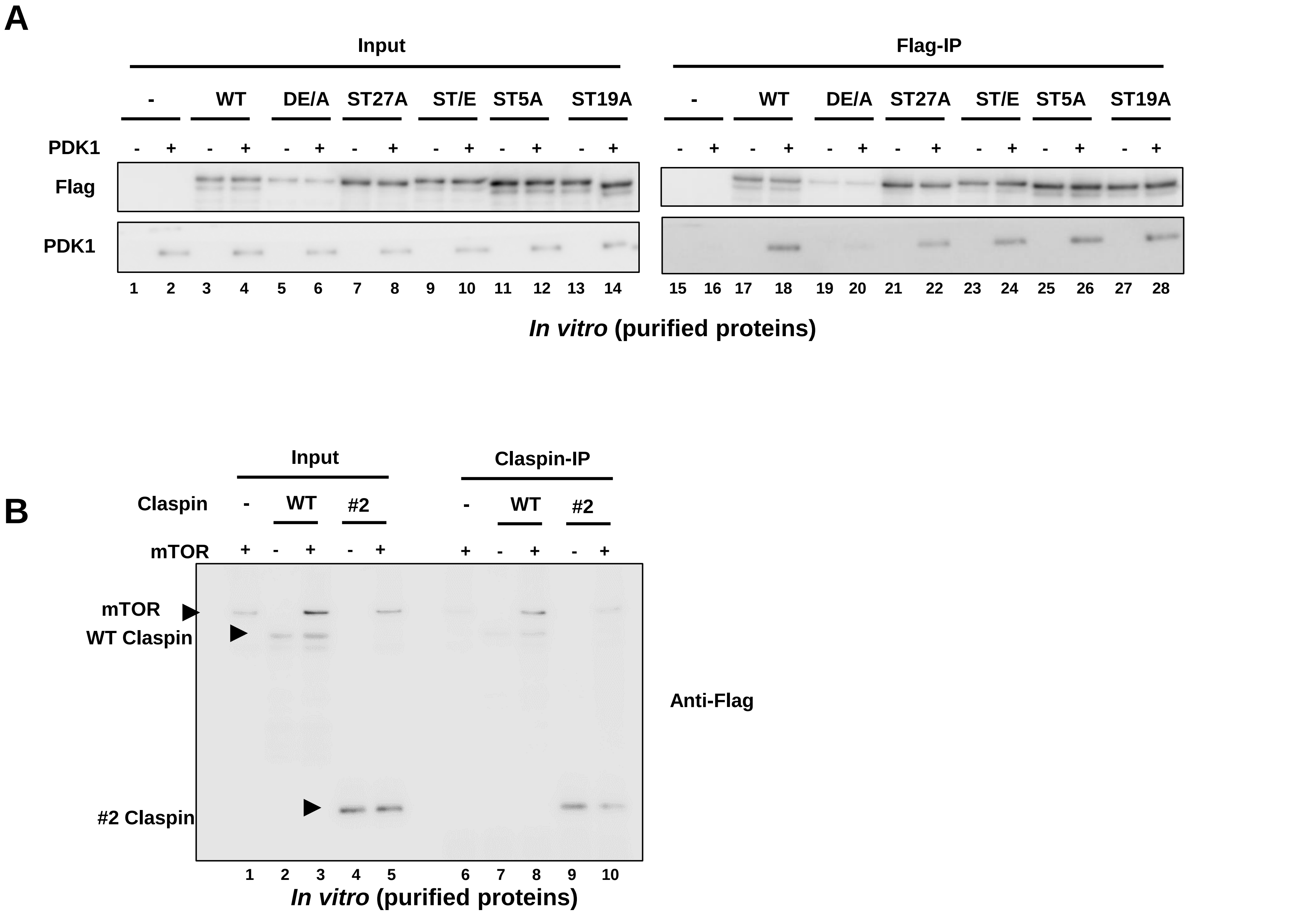
A
Input
Flag-IP
-
WT
DE/A
ST27A
ST/E
ST5A
ST19A
PDK1
Flag
PDK1
1 2 3 4 5 6 7 8 9 10 11 12 13 14
15 16 17 18 19 20 21 22 23 24 25 26 27 28
In vitro (purified proteins)
B
Input
Claspin-IP
Claspin
-
WT
#2
mTOR
mTOR
WT Claspin
#2 Claspin
Anti-Flag
1 2 3 4 5 6 7 8 9 10
In vitro (purified proteins)

Yang_Supplementary Figure S9

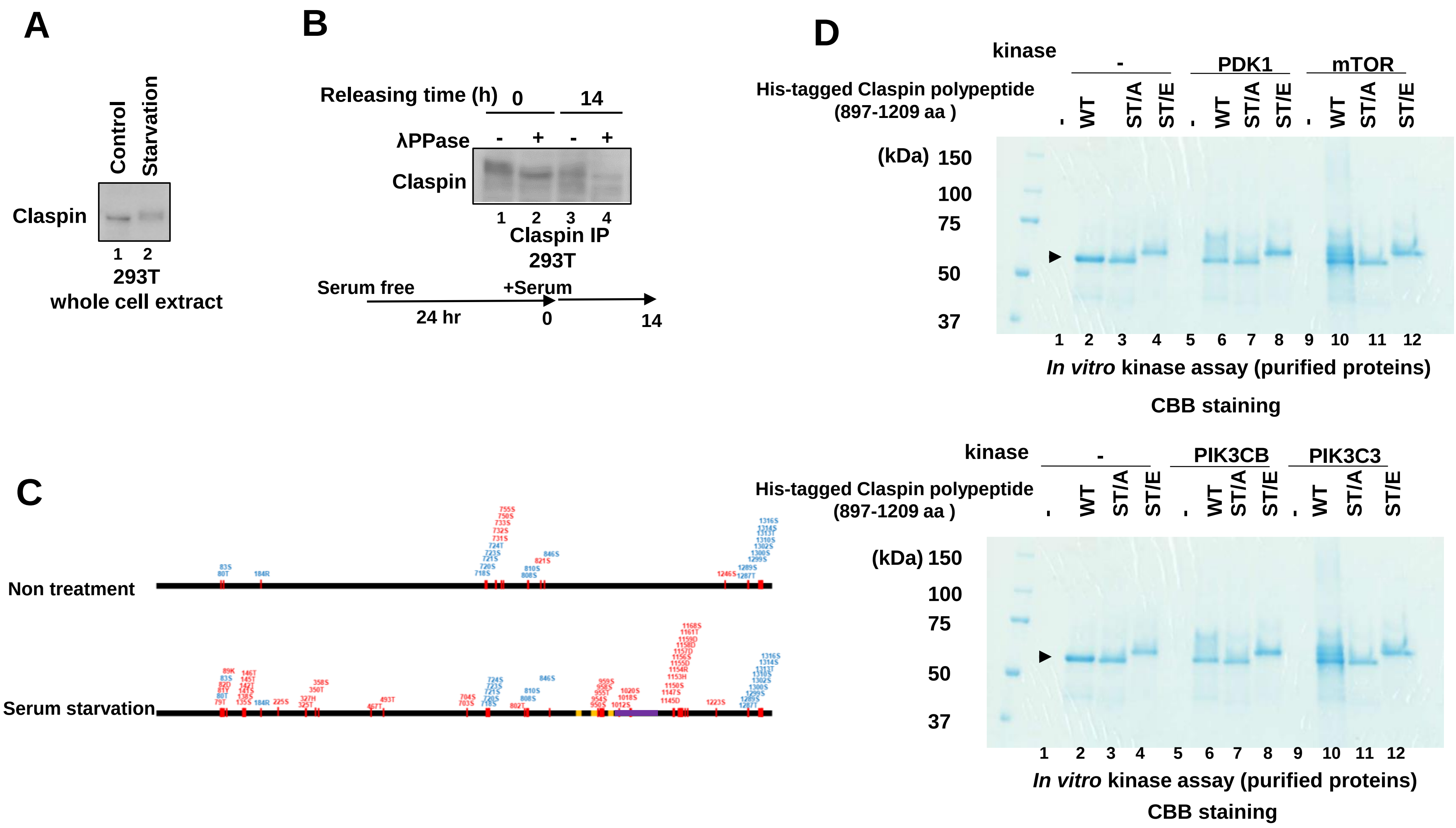

**A**

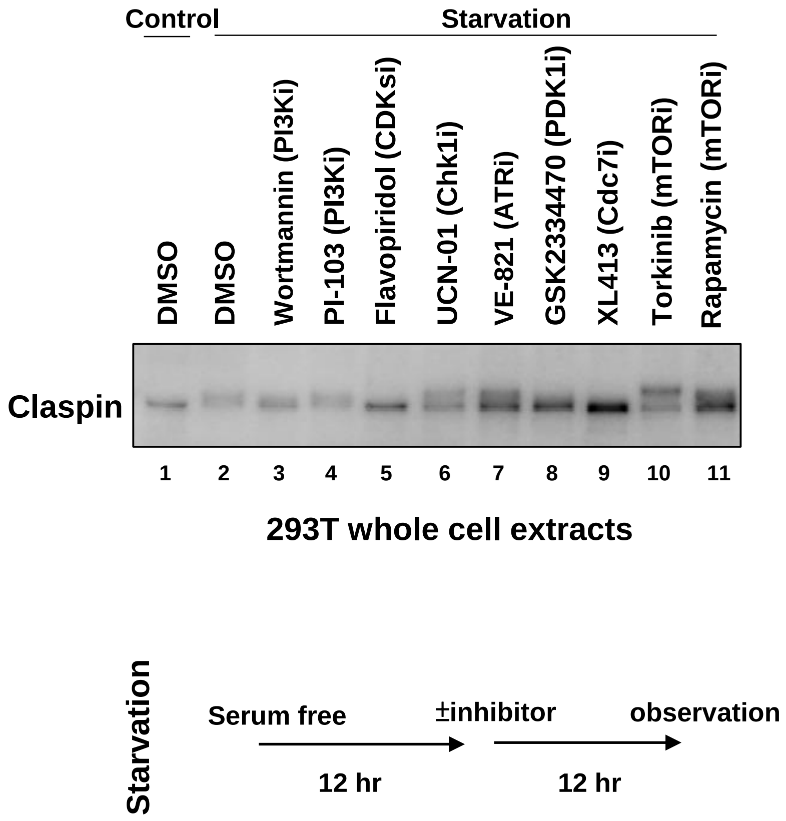
Control
Starvation
DMSO
DMSO
Wortmannin (PI3Ki)
PI-103 (PI3Ki)
Flavopiridol (CDKsi)
UCN-01 (Chk1i)
VE-821 (ATRi)
GSK2334470 (PDK1i)
XL413 (Cdc7i)
Torkinib (mTORi)
Rapamycin (mTORi)
Claspin
1 2 3 4 5 6 7 8 9 10 11
293T whole cell extracts
Starvation
Serum free
±inhibitor
observation
12 hr
12 hr

**B**

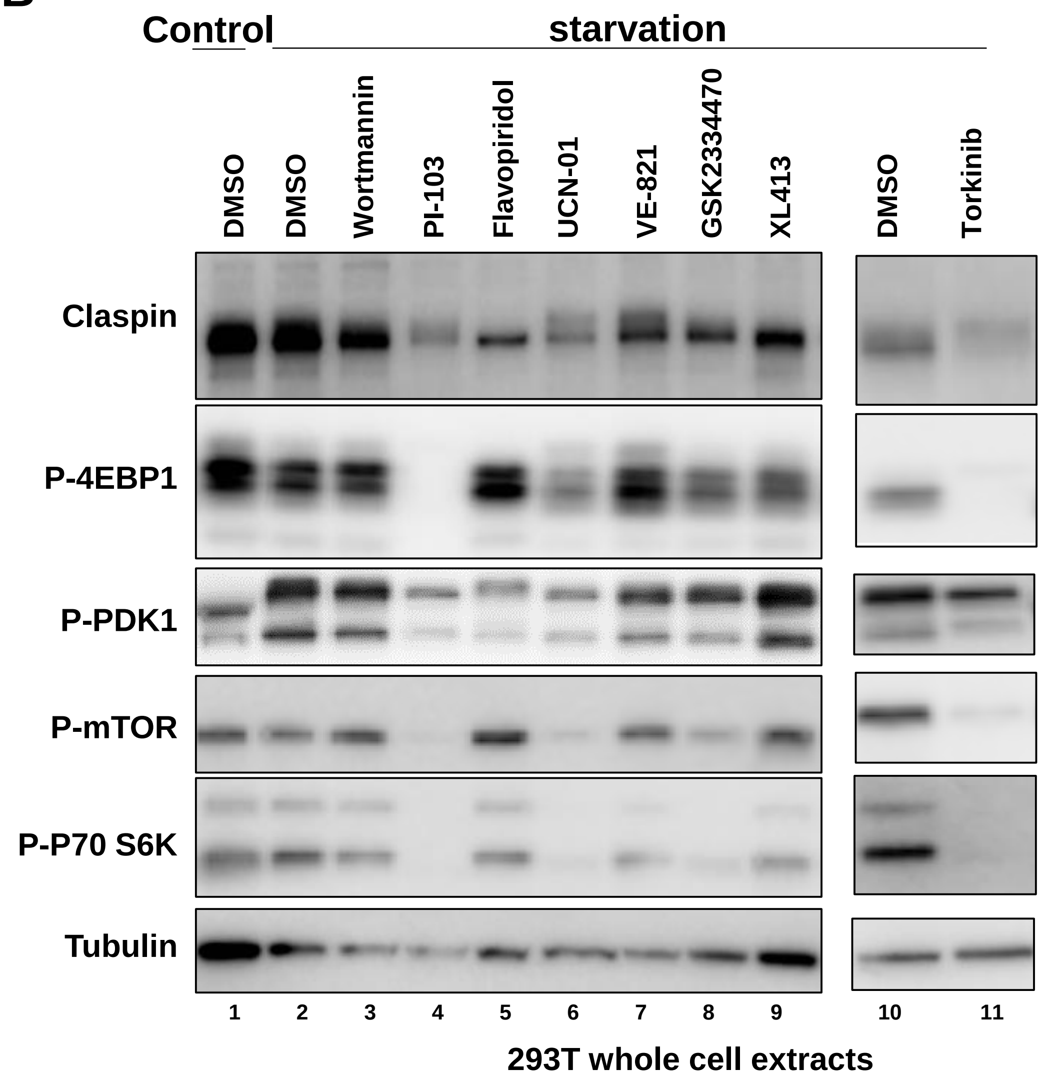
Control
starvation
DMSO
DMSO
Wortmannin
PI-103
Flavopiridol
UCN-01
VE-821
GSK2334470
XL413
DMSO
Torkinib
Claspin
P-4EBP1
P-PDK1
P-mTOR
P-P70 S6K
Tubulin
1 2 3 4 5 6 7 8 9 10 11
293T whole cell extracts

Yang_Supplementary Figure S11

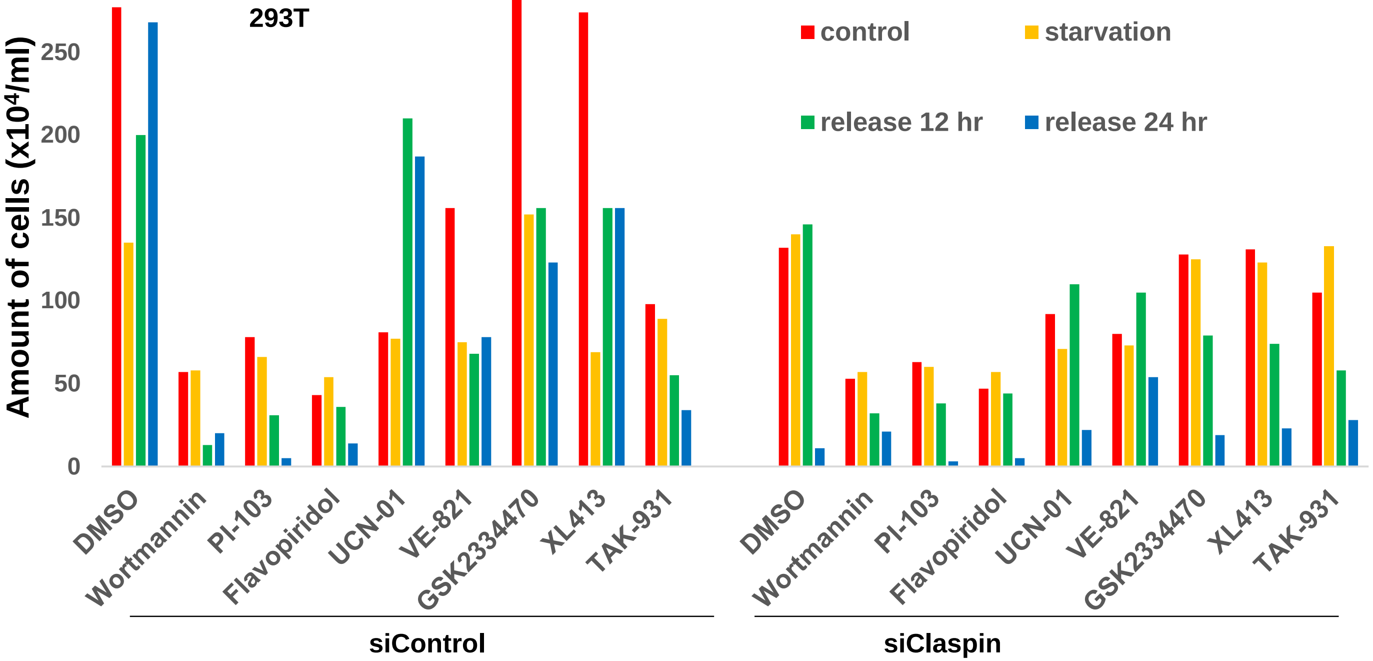


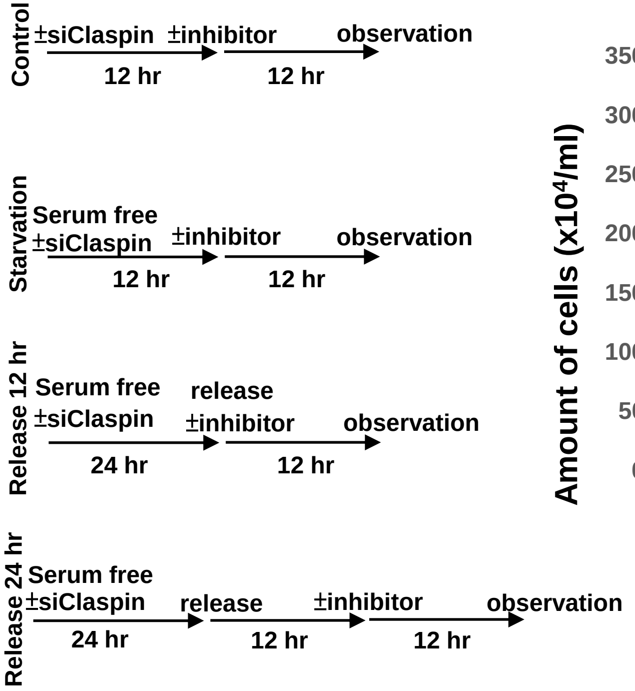


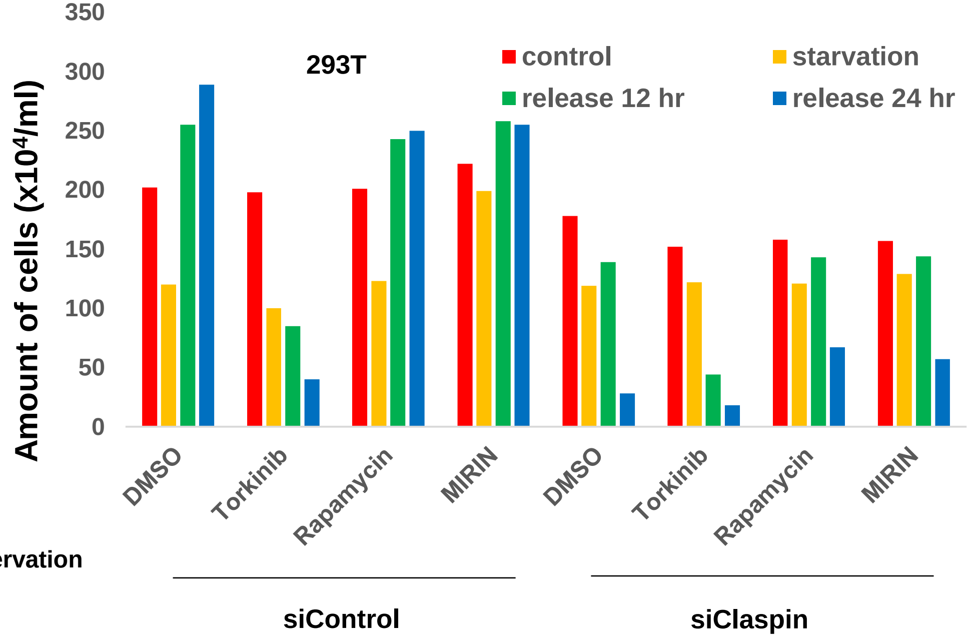

Yang_Supplementary Figure S12

**A**

**MEF *Claspin f/-* vs *Calspin-/-***

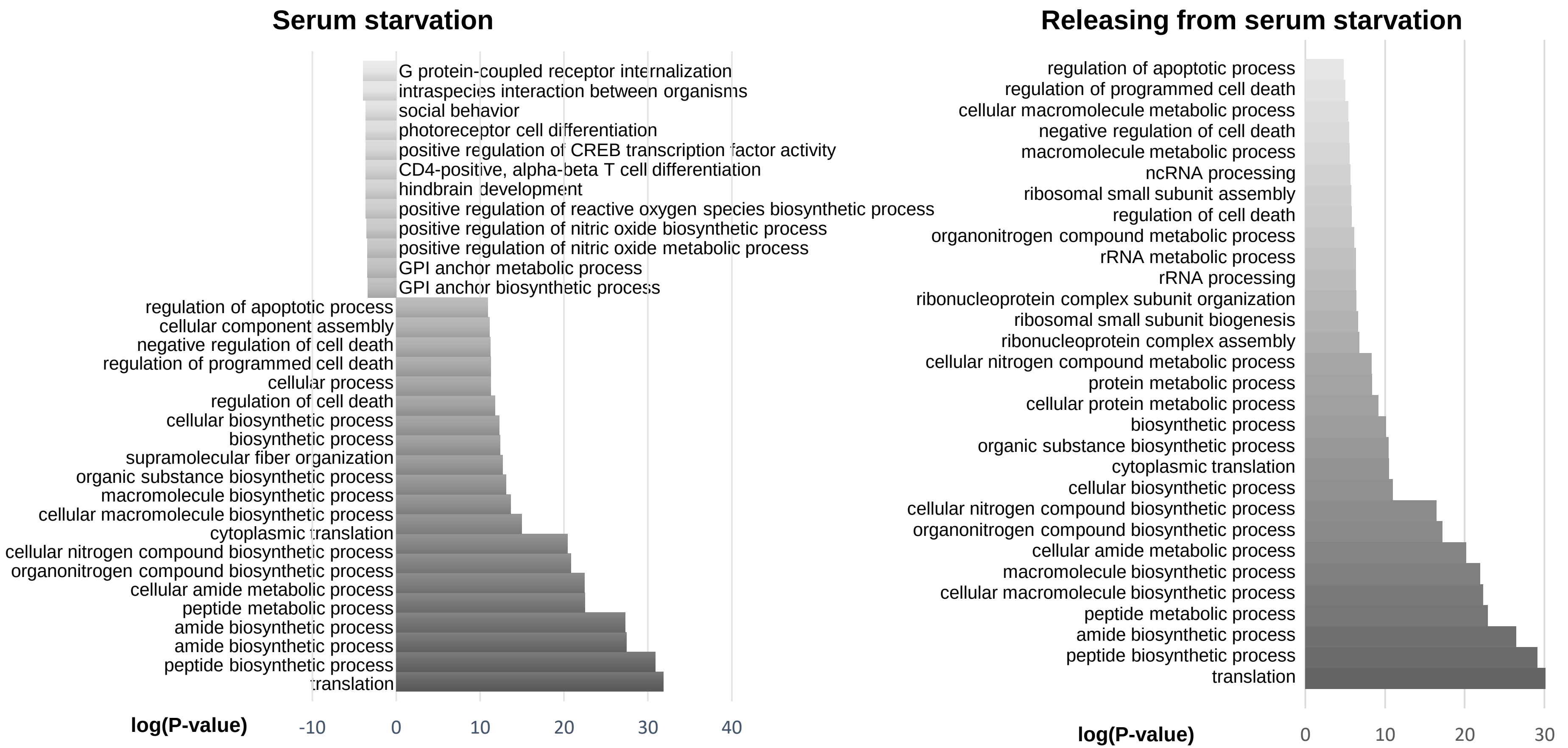


**B**

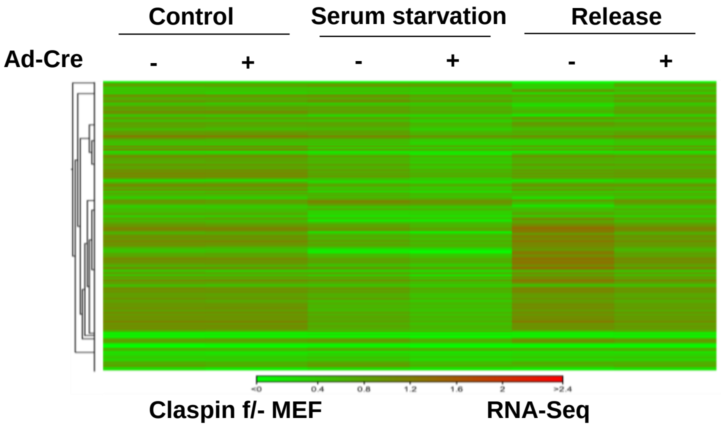
